# Structural insights into cholesterol sensing by the LYCHOS-mTORC1 pathway

**DOI:** 10.1101/2024.11.21.624772

**Authors:** Shang Yu, Jin-hui Ding, Weize Wang, Jia-le Wang, Peng Zuo, Ao Yang, Zonglin Dai, Yuxin Yin, Jin-peng Sun, Ling Liang

## Abstract

The mechanistic target of rapamycin complex 1 (mTORC1) pathway regulates cellular metabolism and growth by coordinating nutrient resources, including cholesterol, and its aberrant activation is linked to various age-related diseases. LYCHOS is a cholesterol sensor on the lysosome and bound to the GATOR1 complex, a GTPase-activating protein for the Rag GTPase, at high cholesterol concentrations, thereby activating the protein kinase mTORC1. However, how LYCHOS senses cholesterol and transduces signal to GATOR1 remain largely unknown. Here we report six cryo-electron microscopy structures of human LYCHOS, depicting five distinct states. These are categorized into a contracted state when complexed with a sufficient amount of the cholesterol analogue cholesteryl hemisuccinate (CHS), and an expanded state when CHS is deficient. The structure forms a homodimer, within each monomer the transmembrane region is divided into a permease-like domain (PLD) and a GPCR-like domain (GLD) with two clearly defined adjacent cholesterol binding sites between them. The PLD shares a conserved Na^+^/H^+^ antiporter (NhaA) fold, which much resembles plant auxin transporter PINs. Cholesterol locates between PLD and GLD and cholesterol binding induces a translation of GLD towards PLD and exposes the cytosolic extension of transmembrane 15, which mediates the interaction between LYCHOS and GATOR1. Strikingly, structure-guided mutations of Gly702 in GLD of LYCHOS increase its affinity for cholesterol, leading to sustained mTORC1 activation in cells. This indicates that LYCHOS’s moderate affinity for cholesterol is crucial as a cholesterol sensor. Our results not only showed a solute carrier mechanistically coordinates a GPCR domain, elucidating the structural mechanism of cholesterol sensing by the mTORC1 pathway on the lysosome; but also provides a structural basis for developing inhibitors that selectively target to mTORC1 pathway to treat age-related diseases by blocking LYCHOS in its expanded state.

## Introduction

The master growth regulator, mechanistic target of rapamycin complex 1 (mTORC1) kinase, is activated at the lysosome. Nutrients, including cholesterol, communicate with mTORC1 via the heterodimeric Rag GTPases (RagA or RagB bound to RagC or RagD)^1–3^. When nutrients are abundant, RagA binds GTP and RagC binds GDP and this complex is active by recruiting mTORC1 to the lysosomal surface^4,5^, where the kinase activity of mTORC1 is stimulated by RHEB (Ras homologue enriched in brain)^6^. Upon nutrient deprivation, the Rag GTPases becomes inhibited by adopting the opposite nucleotide loading state, which cannot bind mTORC1 and leads to cytoplasmic localization of mTORC1^5^. However, the intrinsic GTP hydrolysis rates of the Rag GTPases are slow, making it difficult to quickly change the nucleotide state to adapt to nutrient level change. Two GTPase-activating protein (GAP) complexes, FLCN-FNIP2 (Folliculin-Folliculin Interacting Protein 2) and GATOR1, stimulating GTP hydrolysis of RagC/D and RagA/B, respectively^7–9^, thereby activating and inactivating Rag GTPases, respectively. GATOR1 has three stably interacting subunits: DEPDC5, NPRL2 and NPRL3, with GAP activity residing in the NPRL2 subunit^10,11^. The interaction between GATOR1 and KICSTOR, a complex consisting of four stably interacting subunits: SZT2, KPTN, ITFG2 and C12orf66, facilitates GATOR1-simulated GTP hydrolysis on RagA/B and leads to inactivation of mTORC1^12^.

Cholesterol, as a major component of cell membranes, is vital for proper cellular and systemic functions. Dysregulated cholesterol levels and metabolism are implicated not only cardiovascular disease but also neurodegenerative diseases and cancers^13^. Receptor-mediated endocytosis of low density lipoproteins (LDLs) deliver cholesterol to lysosomes from the extracellular space^14^. Cholesterol on the lysosomal limiting membrane directly participates in the recruitment and activation of mTORC1 in cells and *in vitro*^15^. Oxysterol binding protein (OSBP), a sterol carrier, is found at ER-lysosome membrane contact sites, where it transports cholesterol from the ER to the lysosome, allowing mTORC1 activation^16^. In contrast, the cholesterol transporter Niemann-Pick C1 (NPC1) promotes cholesterol export from the lysosomal surface, inhibiting mTORC1 signaling^15^. How cholesterol interacts with the mTORC1-scaffolding machinery is not well understood. The lysosomal transmembrane protein, SLC38A9 (Solute Carrier Family 38 Member 9), participates in cholesterol dependent activation of mTORC1 through putative sterol-interacting motifs within its transmembrane (TM) 8^15^. However, SLC38A9 has been identified primarily as an arginine sensor that relays arginine abundance to mTORC1^17,18^, and structural evidence for its cholesterol sensing still remains lacking, whereas a dedicated cholesterol sensor has yet to be identified.

Lysosomal cholesterol signaling (LYCHOS, also known as G protein–coupled receptor 155, GPR155, DEP domain containing 3) has recently stood out because it enables cholesterol-dependent activation of mTORC1 independent of amino acid levels^19^. LYCHOS is composed of a 10-TM domain with similarity to solute carriers (SLCs) in the N-terminal; a 7-TM with similarity to GPCRs in the middle, which contains a large loop named LYCHOS effector domain (LED) between TM helices 15 and 16; and a Dishevelled, Egl-10, and Pleckstrin (DEP) domain in the C-terminal region. When cholesterol concentrations on the lysosome membrane are high, LYCHOS disrupts the GATOR1-KICSTOR interaction by its LED and activates mTORC1. The highly conserved residue Tyr551 in LED at the cytosolic side of TM15 is critical for its interaction with GATOR1 as Y551A fails to bind GATOR1 and is unable to activate mTORC1 at high cholesterol concentrations^19^. However, the molecular mechanism of cholesterol sensing by LYCHOS on the limiting lysosome membrane is largely unknown. Here we present cryo-electron microscopy (Cryo-EM) structures of human LYCHOS in complex with cholesteryl hemisuccinate (CHS) and other lipids at different conformational states, revealing in atomic detail the mechanism of cholesterol sensing by the mTORC1 pathway.

## Results

### Structure determination of human LYCHOS

Human LYCHOS were expressed in Sf9 cells and purified through tandem affinity and size exclusion chromatography (SEC) in the absence and presence of cholesterol or CHS. Although all purified LYCHOS proteins, including its mutant Y57A, exhibited good solution behavior (Extended Data Fig. 1a-e), we can only obtain high resolution structures of LYCHOS in the presence of CHS. Microscale thermophoresis (MST) experiments were then carried out with purified LYCHOS with CHS and purified recombinant GATOR1, which show a binding affinity about 7.6 μM, in consistence with the binding affinity between purified MBP-LED^LYCHOS^ and GATOR1 (12 μM) (Fig. 1a). Using single-particle cryo-EM, we determined structures of expanded wild-type LYCHOS (LYCHOS_WT_E_) (Fig. 1b and Extended Data Fig. 2) at a resolution of 3.76 Å, contracted wild-type LYCHOS (LYCHOS_WT_C_) at a resolution of 3.31 Å (Fig. 1c and Extended Data Fig. 2), and contracted LYCHOS_Y57A_ (LYCHOS_Y57A_C_) at a resolution of 3.21 Å (Fig. 1d and Extended Data Fig. 3). 3D variability analysis (3DVA) of the LYCHOS_WT_ map reveal a dynamic swing of GLD around PLD (Supplementary Video 1). The 3D maps are all of high quality in the PLD, enabling us to build accurate model of residues 34-370 (Extended Data Figs. 4, 5). However, the GLD, LED and DEP are all highly dynamic in LYCHOS_WT___C_ and LYCHOS_WT___E,_ therefore we refined these domains to high resolution by focused local refinement and docked these domains predicted by AlphaFold2 separately and adjusted manually by bulky side chain residues (Extended Data Fig. 4). The map of LYCHOS_Y57A_C_ has the best overall quality and its GLD and DEP are clearly resolved (Figs. 1e-f and Extended Data Figs. 4b, 5), therefore we use it in the analysis of the overall structure of LYCHOS.

**Figure 1.**
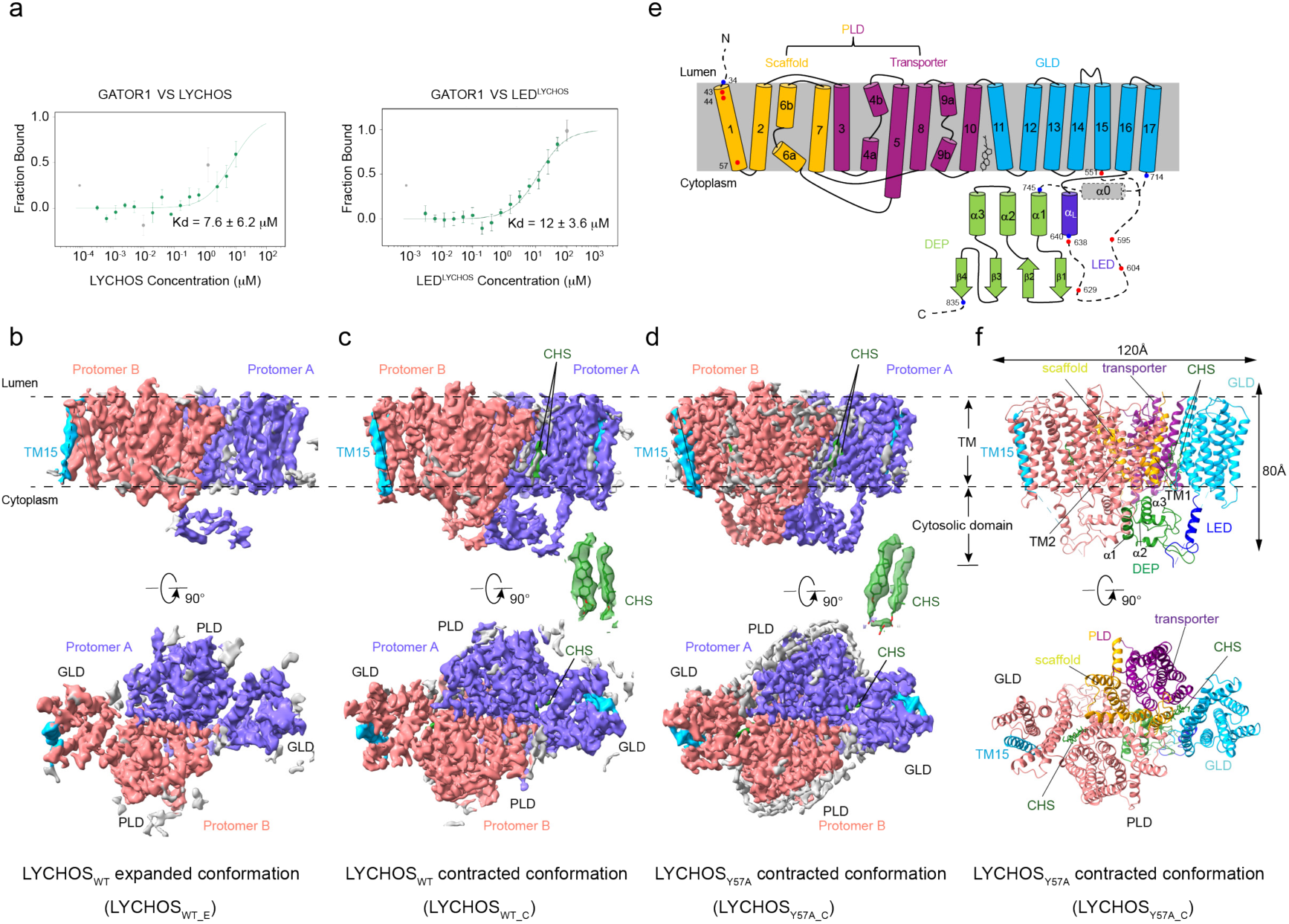
| Cryo-EM structures of human LYCHOS. a, MST experiments showing that our purified LYCHOS with CHS has a similar binding affinity for GATOR1 (Kd = 7.6 μM) compared with LED (Kd = 12 μM). b-d, Cryo-EM maps of lipids bound expanded LYCHOS_WT_E_, CHS bound contracted LYCHOS_WT_C_ and CHS bound contracted LYCHOS_Y57A_. The two protomers are colored in slate blue and light coral and TM15 is shown in deep sky blue. The positions of TM15 of LYCHOS_WT_E_ expanded conformation and LYCHOS_Y57A_C_ contracted conformation are indicated by dashed line and solid line. CHS are shown in green and corresponding cryo-EM map are shown as insets. Other lipid molecules are shown in light gray. e, Schematic of the LYCHOS secondary structure features. Regions that are not visible are shown in dashed lines. Densities for α0 are very poor and also shown in dashed cylinders. Residues that affect LYCHOS function in previous studies are shown as red dots. Resolved boundary residues are shown in blue dots. Scaffold domain, transporter domain, GLD, LED and DEP are shown as gold, purple, deep sky blue, blue and green, respectively. f, Structural model of LYCHOS_Y57A_C_ shown in cartoon representation. Key domains are colored in protomer A as in e. TM15 in protomer B is shown in deep sky blue.

### Overall structure of human LYCHOS

LYCHOS is a symmetric dimer with a twofold rotation axis perpendicular to the membrane plane and a distinct concavity extending into the membrane from the side of lysosomal lumen along this axis (Extended Data Fig. 6a). Each monomer contains 17 transmembrane helices (TM) with TM1–TM10 constituting the PLD and TM11–TM17 constituting the GLD (Fig. 1e). The overall fold of the PLD is similar to that of auxin transporters^20–22^, with a root mean square deviation (RMSD) of 2.5 Å over 310 Cα atoms (DALI Z-score: 29.2) to Arabidopsis thaliana PIN1 (PDB: 7Y9V) when they are aligned (Extended Data Fig. 6b). The GLD is similar to class F GPCR, with an RMSD of 3.2 Å over 265 Cα atoms (DALI Z-score: 21.9) to Frizzled4 receptor (PDB: 6BD4) when they are aligned (Extended Data Fig. 6c). Like PINs, PLD’s ten transmembrane helices (TM1–TM10) comprise an inverted repeat of five transmembrane helices (Extended Data Fig. 6d). In each repeat, the fourth helix is bent by a conserved serine or a conserved proline residue in the middle: Ser144 in TM4 and Pro324 in TM9, identical to Pro111 in TM4 and Pro579 in TM9 of PIN1, respectively (Extended Data Figs. 5, 6e, 6f). These disrupted helices form an X-shaped crossover, indicating a putative unknown ligand binding pocket identical to auxin binding site in PIN1. Like PIN1, the PLD of LYCHOS monomer is divided into two domains: the scaffold domain and the transporter domain (Figs. 1e, 1f and Extended Data Fig. 6f). The scaffold domain comprises helices TM1, TM2, TM6 and TM7 and creates a large interface (1870 Å^2^) to the other monomer in the dimeric complex. This interface is primarily mediated by TM2 and TM7, with lipids in a groove between TM1, TM11 and the kinked TM6 (Figs. 1b-d and Extended Data Fig. 6g, h). Another lipid with an aliphatic tail was observed by sticking into the pocket constituted by TM3 and TM8 (Extended Data Fig. 6h). The identities of these lipid-like densities remain to be further investigated. The transporter domain consists of helices TM3–TM5 and TM8–TM10 and harbours the central X-shaped crossover.

DEP also forms a dimer interface involving α1 and α2 (Fig. 1f and Extended Data Fig. 6i). The α0 connecting GLD and DEP is highly dynamic. The loop connecting α2 and α3 interacts with the loop connecting TM1 and TM2 (Fig. 1f and Extended Data Fig. 6i). Interestingly, although the interaction details cannot be unambiguously assigned, we can see LED interacts with DEP (Fig. 1f and Extended Data Fig. 6i).

### Recognition of cholesterol by LYCHOS

The ability for the interaction between our purified LYCHOS and GATOR1 indicates LYCHOS_WT_ binds cholesterol or at least its analogue CHS. There are several densities with unknown identities in the cleft between PLD and GLD. The highly dynamic feature of GLD hampered accurately model cholesterol binding site. After 3D classification, we acquired an asymmetric class in which one of the two protomers shows clear cholesterol-shaped densities with an overall resolution of 3.31 Å (Extended Data Figs. 2, 4a). Interestingly, we found two well-defined patches of density with a cholesterol shape in the potential cholesterol-binding cleft (CBC) between PLD and GLD (Fig. 2a and Extended Data Fig. 4a) and the local resolution around cholesterol-binding sites is higher than 3.3 Å (Extended Data Fig. 2f). We name them as cholesterol binding sites CBS1 and CBS2 in the cytosolic side near TM1 and TM10, respectively (Fig. 2b). Both CBS1 and CBS2 had EM densities that could be unambiguously fitted with CHS (Extended Data Fig. 4a). Overall, the pocket is enclosed by TMs 1/10/11/16/17. CBS1 is composed of Phe43, Phe50, Leu54, Tyr57 of TM1, Phe352, Tyr359 of TM10, Ile386 of TM11 and Phe705, Phe708 of TM17 (Fig. 2c). The aromatic ring of Try57 makes a π-anion interaction with the hydrophilic carboxyl head group of CHS and forms π-π stacking with the tetracyclic ring. Phe50 and Phe705, Phe352 and Phe708 form π-π stacking with the tetracyclic ring. The hydrophobic tail is inserted into a pocket formed by Phe43, Phe50, Tyr359 and Ile386. Overall, the CBS1 cavity is primarily formed by TM1 and TM10 of PLD and subsequently sequestered by TM17 of GLD in the contracted state.

**Figure 2.**
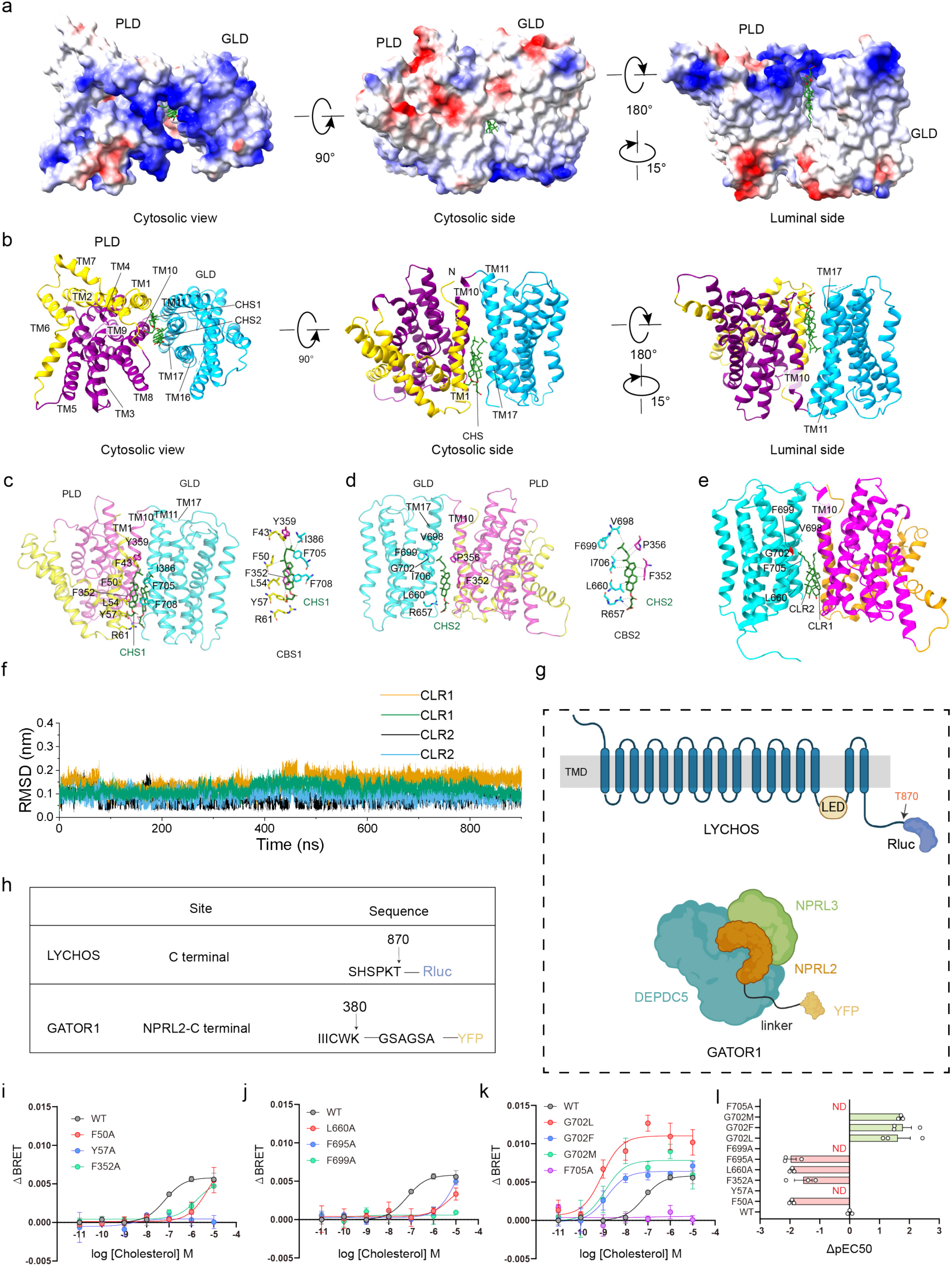
| Cholesterol binding sites. a, Electrostatic potential surfaces of protomer A of the LYCHOS_WT_C_ structure. CHS molecules are shown as sticks. b, Cartoon representation of the LYCHOS_WT_C_ structure. CHS molecules are shown as sticks. c, Details of the cholesterol binding site CBS1 of LYCHOS_WT_C_. CHS molecules are shown in green and sticks. Hydrophobic residues interacting with CBS1 are shown in sticks. Scaffold domain (PLD), transporter domain (PLD) and GLD are shown as gold, purple and deep sky blue, respectively. d, Details of the cholesterol binding site CBS2. CHS2 and hydrophobic residues interacting with CBS2 are shown in sticks. Cartoon model for TM16 and TM11 are omitted for convenience to read. L660 from TM16 is shown in stick. e, The molecular interaction between cholesterol molecules and LYCHOS. The interactions were revealed by 900 ns molecular dynamics simulation. Cholesterol (CLR) molecules are shown in green and sticks. Hydrophobic residues interacting with CBS1 are shown in sticks. Scaffold domain (PLD), transporter domain (PLD) and GLD are shown as gold, purple and deep sky blue, respectively. Gly702 is shown in red. f, The root mean square deviation (RMSD) of the cholesterol molecules in two CBSs during the 900 ns simulation for two trajectories. g, Schematic of LYCHOS and GATOR1 modification design, fusing Rluc to the LYCHOS-C-terminus and linking YFP to the NPRL2-C-terminus of the GATOR1 heterotrimer via linker. In response to cholesterol stimulation, LYCHOS recruits GATOR1, and Rluc is brought into proximity with YFP, and energy resonance transfer occurs. h, Detailed description of the Rluc and YFP fusion sites. Rluc is indicated in blue and YFP is indicated in yellow. i-k, Cholesterol dose-dependent profile of LYCHOS recruitment to GATOR1 after CBS mutations as determined by BRET. Data from three independent experiment (n = 3). l, Effect of CBS mutations on cholesterol-activated LYCHOS recruitment of GATOR1. Data from three independent experiment (n = 3). ***P < 0.001; **P < 0.01; *P < 0.05; ND, no detectable signal; ns, no significant difference. Values are shown as the mean±s.e.m. from three independent experiments performed in triplicate. And data were statistically analyzed using one-way ANOVA with Dunnett’s post hoc test.

The TM16, TM17 and TM10 form the second binding site, CBS2, which is adjacent to CBS1 (Fig. 2c,d). This site is composed of Phe352, Pro356 on TM10, Leu660 on TM16 and Val698, Phe699, Phe705, Phe708 on TM17 (Fig. 2d). Discontinuity of TM17 makes sufficient space for the tail of cholesterol to fit inside (Fig. 2d). Gly702 and Gly704 are responsible for this unwinding and are highly conserved (Extended Data Fig. 5).

As both CBS binds CHS, we wondered whether cholesterol binds LYCHOS in the same way as CHS. We substituted CHS with cholesterol and performed molecular dynamics simulations in bilayer phospholipids over the course of 900 ns. As shown in Fig. 2e, cholesterol molecules bind CBS in the same way as CHS despite slightly dynamic, which indicates its weaker binding affinity with LYCHOS when compared with CHS.

To investigate whether the two cholesterol binding sites are important for LYCHOS to sense cholesterol concentrations, we developed a Renilla luciferase (Rluc)/YFP bioluminescence resonance energy transfer (BRET) method to monitor the recruitment of GATOR1 components after LYCHOS binds cholesterol. Rluc was fused to the C-terminus of LYCHOS, while YFP was incorporated into the C terminal of NPRL2 in the components of GATOR1 (Fig. 2g-h). By LYCHOS-Rluc/NPRL2-YFP recruiting experiments, we found that the cholesterol promoted recruiting NPRL2 (GATOR1) to LYCHOS in a concentration dependent manner, with an EC50 of 165.1 nM, which is in consistent with previous report^19^. Moreover, F50A, Y57A, F352A, L660A, F695A, F699A or F705A mutations in these two CBS sites significantly reduced LYCHOS activation in response to cholesterol stimulation (Fig. 2i-l), which supports our structural findings. Interestingly, the G702L, G702M and G702F mutations significantly enhanced cholesterol-induced GATOR1 recruitment to LYCHOS (Fig. 2k-l). Structural analysis indicated that the relatively bulky side chains of leucine and phenylalanine enhance hydrophobic interactions with the hydrophobic tail of cholesterol. We further examined the effects of Gly702 mutations G702L, G702M and G702F on mTORC1 activity *in vivo*. As expected, all mutations resulted in constitutive activation of mTORC1 (Extended Data Fig. 7). Taken together, our data support the CBC between PLD and GLD acts as the signal center for cholesterol sensing and both CBS1 and CBS2 are critical for cholesterol sensing.

### LYCHOS adopts an expanded conformation without sufficient cholesterol

As a cholesterol sensor rather than a mere cholesterol-binding protein, LYCHOS crucially undergoes conformational changes upon cholesterol binding. We observed an expanded state of LYCHOS (LYCHOS_WT_E_) distinct from the previously discussed contracted state. While the PLD and GLD maintain overall similarity to the contracted state (RMSD of 0.52 Å over 297 Cα atoms and 1.16 Å over 191 Cα atoms, respectively), the GLD in LYCHOS_WT_E_ adopts a unique conformation, expanding away from the PLD (Fig. 3a). Comparing the expanded and contracted states reveals a rigid-body rotation of the GLD towards the PLD in the contracted state, pivoting around Leu375 on the non-cytosolic side of TM11 (Fig. 3b,c and Supplementary Video 2). Among GLD’s TMs, TM15’s cytosolic side exhibits the most significant shift (Fig. 3c), with the crucial GATOR1-binding residue, Tyr551, positioned at its cytosolic extension. However, the cytosolic extension of TM15 is not visible in our cryo-EM map. To address this, we incorporated this region using an AlphaFold-predicted model, aligning it with the visible portion of TM15 in our structure. In LYCHOS_WT_E_, Tyr551 is located closer to the membrane compared to LYCHOS_WT_C_. The distances between Tyr551 and the detergent micelle density are 3 Å and 7.5 Å for LYCHOS_WT_E_ and LYCHOS_WT_C_, respectively (Fig. 3d, e). Molecular dynamics simulations in bilayer phospholipids over 1000 ns corroborate these structural observations: Tyr551 is nearly buried in the membrane in the expanded state but exposed to the cytoplasm in the contracted state (Fig. 3f, g). These findings suggest that in cholesterol-abundant environments, cholesterol molecules act as molecular glues, inducing LYCHOS to adopt a contracted conformation. This conformational shift brings the GLD closer to the PLD, consequently exposing Tyr551 and promoting LYCHOS-GATOR1 interaction. Conversely, in the expanded state, Tyr551 is nearly buried and unable to bind GATOR1. The critical nature of Tyr551 for LYCHOS-GATOR1 interaction is further evidenced by the failure of LED-Y551A to bind GATOR1.

**Figure 3.**
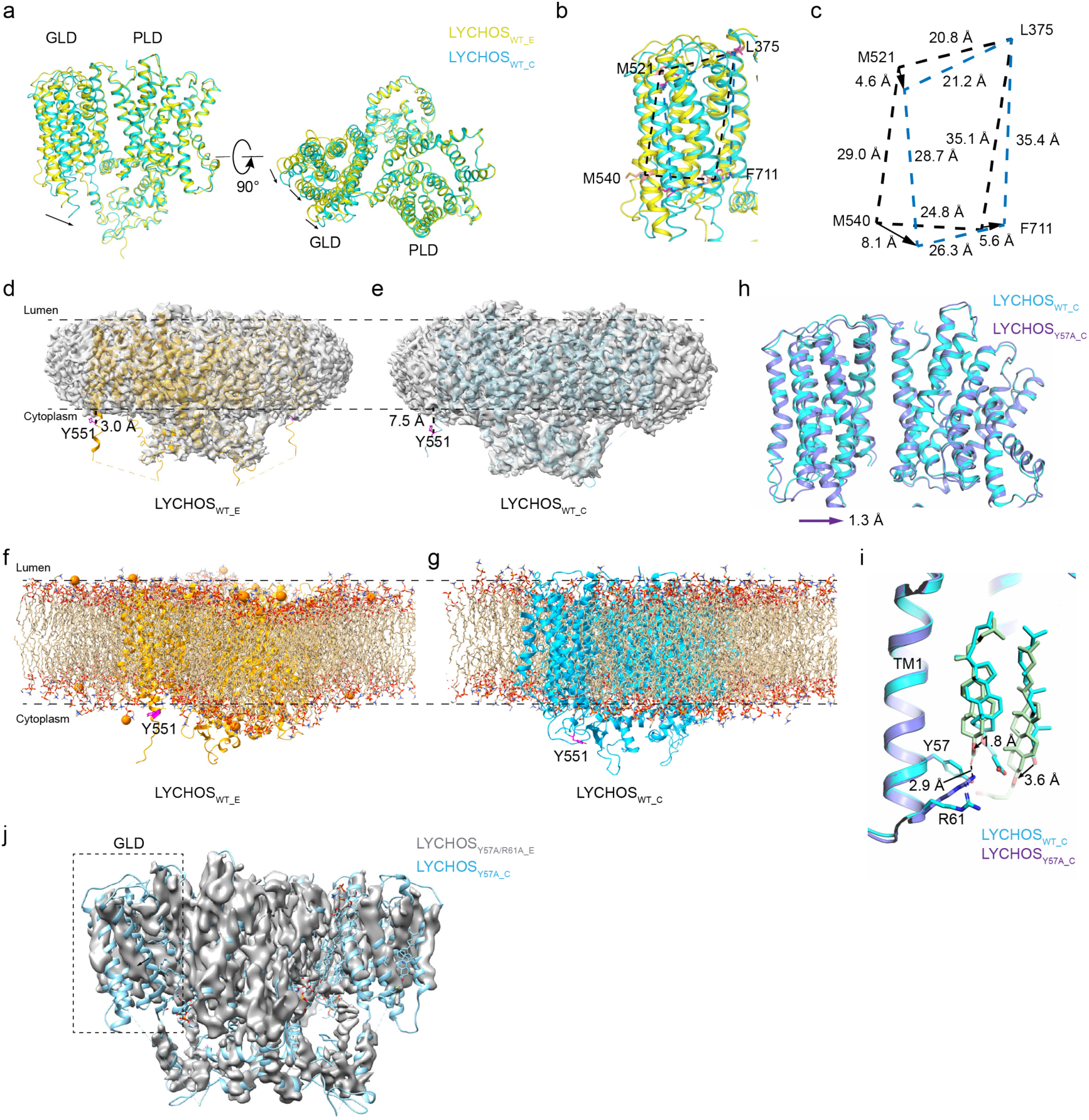
| LYCHOS adopts an expanded conformation without sufficient cholesterol. a, Cartoon representation of LYCHOS_WT_C_ (cyan) and LYCHOS_WT_E_ (yellow), superimposed based on the PLD domain of chain A in LYCHOS_WT_C_ and chain B in LYCHOS_WT_E_. b, Conformational displacements of the GLD relative to the PLD between LYCHOS_WT_C_ (cyan) and LYCHOS_WT_E_ (yellow). Cα atoms of residues used to measure displacements are shown in sticks and labeled. c, Schematic representation of the displacement of the four residues highlighted in b. Dashed lines indicate distances between the Cα atoms within each state. Solid arrows indicate the absolute distance between Cα atoms of each residue between LYCHOS_WT_E_ and LYCHOS_WT_C_. d,e, AlphaFold model are fitted based on the TM15 and the distance between the Cα atom of Tyr551 and detergent micelle are shown in dashed lines and labeled. f,g, The distance between Tyr551 and bilayer phospholipids is larger in LYCHOS_WT_C_ when compared with LYCHOS_WT_E_. The bilayer phospholipids are shown in stick. Tyr551 is shown in stick and colored in magenta. The states were revealed by 1 μs molecular dynamics simulation. h, Cartoon representation of LYCHOS_WT_C_ (cyan) and LYCHOS_Y57A_C_ (slate), superimposed based on the PLD domain. i, Conformational displacements of the LYCHOS_WT_C_ (cyan) bound CHS molecules (cyan) and LYCHOS_Y57A_C_ (slate) bound CHS molecules (green) relative to the PLD. Solid arrows indicate the absolute distance between carboxylate oxygen atoms of CHS molecules between LYCHOS_WT_C_ and LYCHOS_Y57A_C_. j, Superimposition of the LYCHOS_Y57A_C_ structure with the map of LYCHOS_Y57A/R61A_E_. The solid arrow indicates the translation of GLD.

Since Tyr57 is critical for cholesterol sensing, we wondered whether an alanine mutation of this residue would break the interaction between LYCHOS and cholesterol and make LYCHOS in the expanded state. Unexpectedly, although LYCHOS_Y57A_C_ does not bind cholesterol, we can see clear densities in the CBC which resemble CHS, therefore we modeled these densities as CHS (Extended Data Fig. 3 and 4b). We found there exist two CHS molecules in the CBC, with which GLD is locked in contracted state and expose the cytosolic extension of TM15. When compared with LYCHOS_WT_C_, LYCHOS_Y57A_C_ adopts a more contracted conformation with a translation of 1.3 Å towards PLD for the Cα atoms of Met542 and the carboxylate oxygen of CHS transits 1.8 Å due to the loss of the bulky side chain of Tyr57 (Fig. 3h). Although alanine mutation of tyrosine disturbs the interaction between CHS and residues 57, CHS forms a salt bridge with Arg61 instead (Fig. 3i). This interaction requires the longer chain of the polar head of CHS, which may explain why CHS still interacts with LYCHOS_Y57A_ while cholesterol cannot. To verify this, we purified the double mutant LYCHOS_Y57A/R61A_, which abolished its interaction with CHS, and solved its structure at a resolution of 3.73 Å (Extended Data Fig. 8a-f). The structure revealed an expanded state (Fig. 3j). These results collectively indicate that LYCHOS is in an expanded state without cholesterol or its analogs in its CBS.

Intriguingly, our purified LYCHOS_Y57A_ revealed an additional conformation beyond LYCHOS_Y57A_C_: an expanded homodimer mediated by kinked TM6, which we designated LYCHOS_Y57A_TM6_E_ (Extended Data Fig. 9). We resolved this structure to a resolution of 4.49 Å. Although side chain details were not discernible, the LYCHOS_Y57A_TM6_E_ homodimer clearly showed TMs of both PLD and GLD. Notably, the structure of each protomer diverged from the contracted state of LYCHOS, instead resembling its expanded state (Extended Data Fig. 9g). These findings suggest that LYCHOS can adopt various conformations depending on environmental conditions, potentially enabling diverse functions. This conformational flexibility may be modulated by lipid interactions, highlighting the complex interplay between protein structure and lipid environment in LYCHOS function.

### GLD and lipids in the dimer interface is essential for cholesterol binding

Previous studies identified the cholesterol recognition amino acid consensus (CRAC) motif on TM1 in PLD as responsible for cholesterol binding^19,23^. To investigate whether PLD alone is sufficient for this interaction, we purified LYCHOS_PLD_ and resolved its structure to 2.73 Å (Extended Data Fig. 10a-f). LYCHOS_PLD_ forms a stable homodimer without cholesterol-like densities around TM1. Instead, we observed numerous phospholipid-like densities at the dimer interface (Fig. 4a and Extended Data Fig. 10). Similar lipid-shaped densities were also evident in LYCHOS_Y57A_C_, positioned between the kinked TM6 of one protomer and TM11 of the other (Fig. 4b). This hydrophobic pocket is surrounded by TM6, TM2 and TM7 of one protomer and TM1, TM11 and TM17 of the other protomer. The densities resemble phosphatidylserine (PS) and therefore we assume these densities as PS for convenience to describe, although the exact identities require further investigation (Fig. 4c). The hydrophobic tail of PS2 sticks deep into the pocket, making hydrophobic contact with CHS in CBS1 (Fig. 4d). Residues Leu47, Leu54, Cys55, Ile58 on TM1, Ile392, Trp396 on TM11 and Phe705, Phe708 on TM17 of one protomer, residues Phe80, Pro83, Phe87 on TM2, Phe221, Met222, Phe229 on TM6 and Pro238, Val241, Phe244 on TM7 of the other protomer make hydrophobic interaction with the lipids (Fig. 4d). Additionally, the positive guanidine group of Arg79 makes electrostatic interaction with the negative phosphate group of PS (Fig. 4d). These interfacial lipids likely play a dual role: stabilizing LYCHOS dimerization and restricting GLD mobility. Their strategic location at both the dimer interface and the PLD-GLD junction suggests a crucial structural and functional significance. This is further evidenced by the non-canonical LYCHOS_Y57A_TM6_E_ homodimer, which lacks these interfacial lipids and consequently exhibits a markedly expanded conformation (Fig. 4e and Extended Data Fig. 9g).

**Figure 4.**
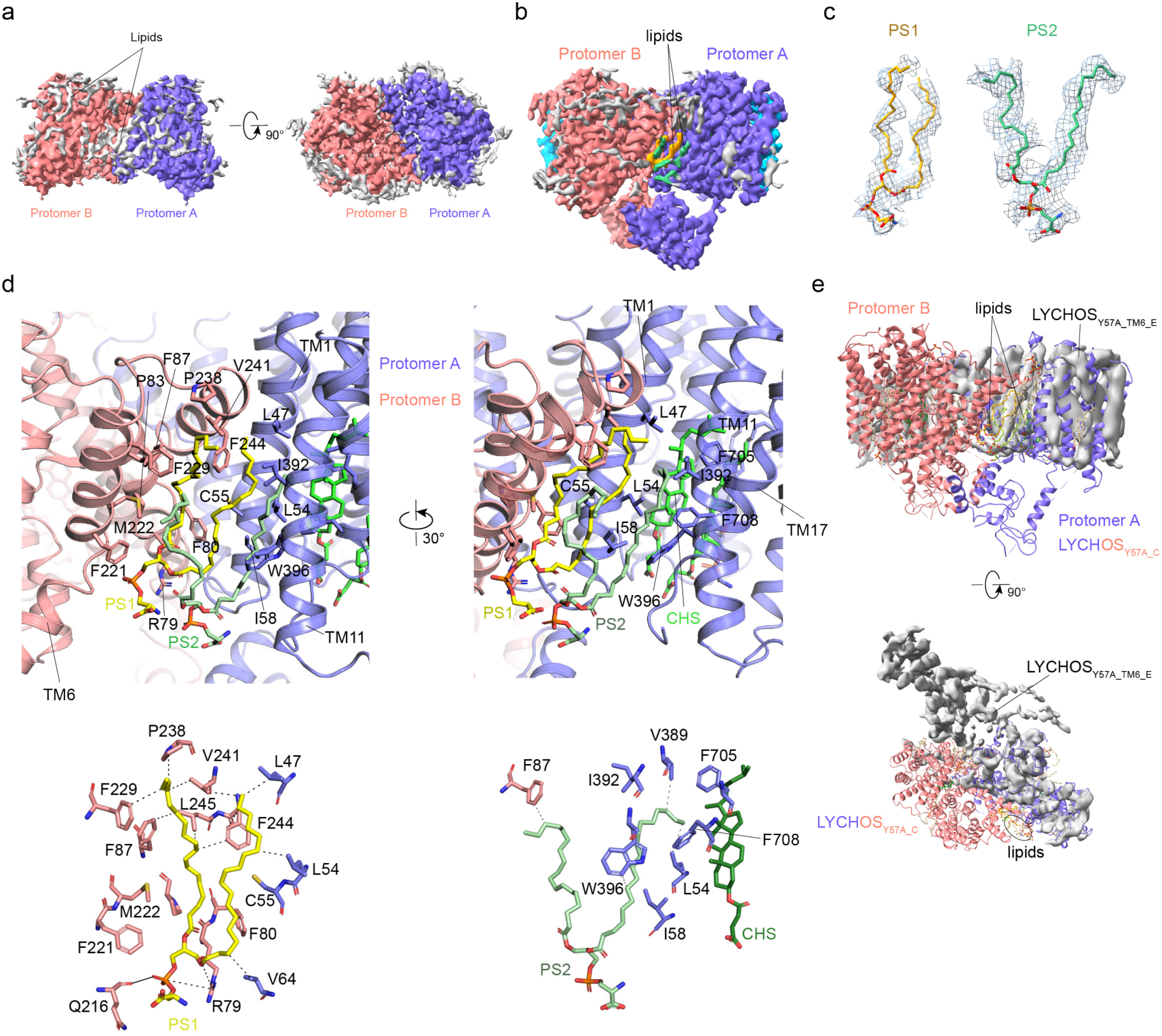
| Lipids reside at the dimerization interface of LYCHOS. a, Cryo-EM maps of LYCHOS_PLD_ homodimer bound with lipids. The two protomers are colored in slate blue and light coral. Lipid molecules are shown in light gray. Two lipid molecules are colored in orange and green, respectively. b, Cryo-EM maps of LYCHOS_Y57A_C_ bound with PS. The two protomers are colored in slate blue and light coral. c, Cryo-EM density maps of PS in the LYCHOS_Y57A_C_ state are shown in blue meshes. d, Details of the interaction between lipids and LYCHOS. PS and CHS molecules are shown as sticks. Residues interacting with lipids are shown in sticks. e, Superimposition of the LYCHOS_Y57A_C_ structure with the map of LYCHOS_Y57A_TM6_E_. The two protomers are colored in slate blue and light coral for LYCHOS_Y57A_C_ and lipids are shown in sticks. The lipids observed in the canonical dimer interface of LYCHOS_Y57A_C_ could not be seen in LYCHOS_Y57A_TM6_E_.

### Working mode for LYCHOS

On the basis of the data available, we propose the following model for cholesterol sensing and signal transducing by LYCHOS (Fig. 5): at low cholesterol concentrations, the cleft between PLD and GLD is occupied by other lipids and GLD adopts an expanded state with Tyr551 buried in the membrane. When cholesterol is sufficient, cholesterol molecules bind in the cleft between PLD and GLD, LYCHOS transits to the contracted state, the GLD rotates and the GATOR1 binding site in the cytosolic extension of TM15 is translated away from the membrane, leading to its interaction with GATOR1, thereby activating the protein kinase mTORC1.

**Figure 5.**
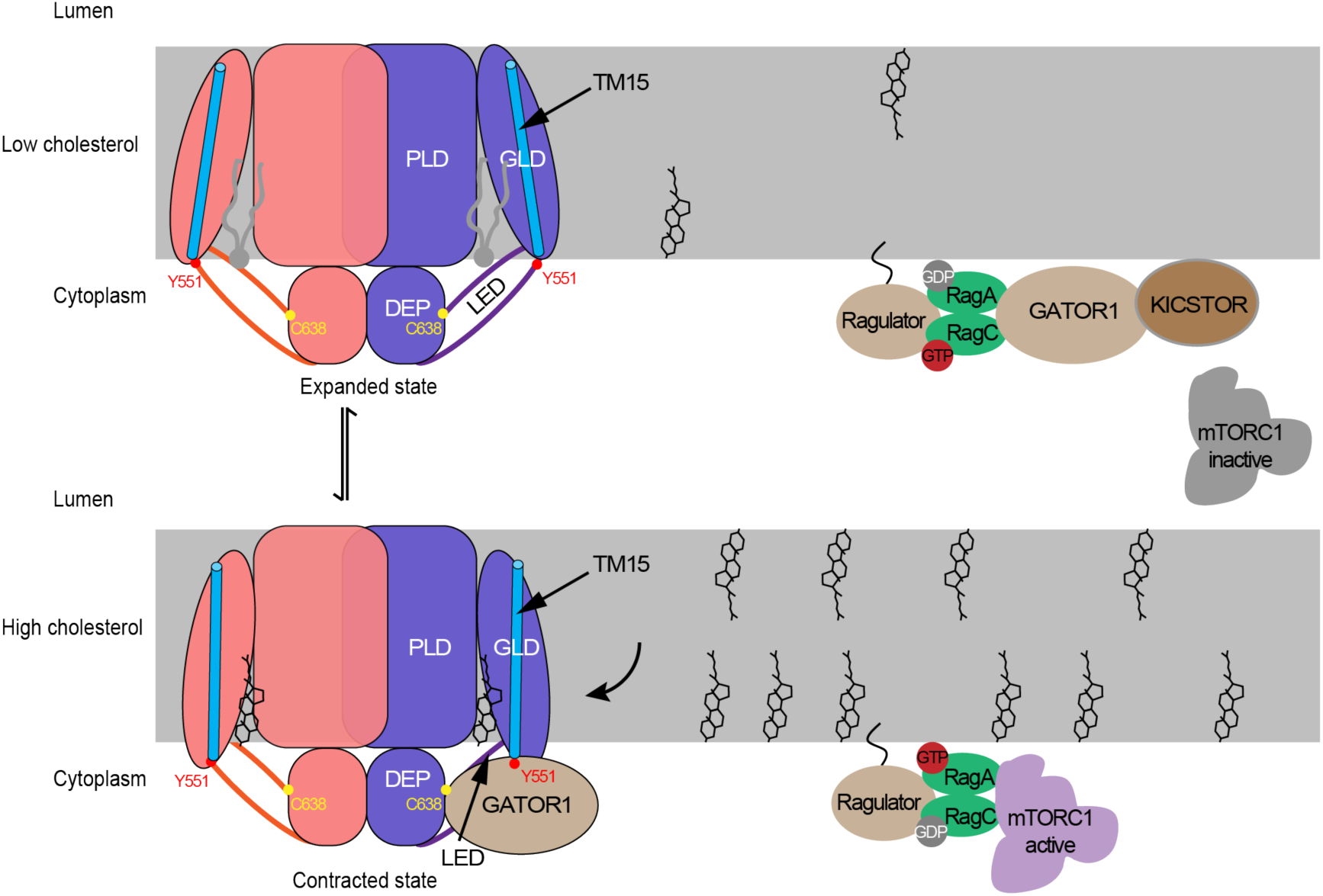
| Proposed mechanism of LYCHOS. Up: Under low cholesterol, LYCHOS is in an expanded state in the absence of cholesterol. GATOR1 promotes GTP hydrolysis of RagA/B, maintaining mTORC1 in the cytosol and inactive. Down: When cholesterol is sufficient, binding of cholesterol induces the contraction of the GLD and the descending of TM15, makes the LED suitable for interaction with GATOR1, thus facilitating the GTP-loaded state of RagA/B that recruits mTORC1 to the lysosome and activate it.

## Discussion

Although recent structural studies of LYCHOS have been conducted^23^, its cholesterol-sensing and signal transduction mechanism remains unclear. Understanding how LYCHOS recognizes cholesterol, changes conformation, and recruits GATOR1 could significantly impact our knowledge of metabolic regulation. LYCHOS’s highly dynamic nature and cholesterol sensing mechanism have been described here. The dynamic GLD makes it suitable for sensing concentration changes of cholesterol. Only when cholesterol concentrations are sufficiently high, binding with LYCHOS will change the conformation of GLD and transduce signal. Meanwhile, our structures reveal at least three states: a canonical contracted homodimer mediated by the TM2/7 as PINs, a canonical expanded homodimer mediated by the TM2/7 and a non-canonical expanded homodimer mediated by the kinked TM6. The contracted state is only seen in the canonical homodimer, whereas the expanded state is seen in both canonical and non-canonical homodimer. It is unknown whether contracted states exist in non-canonical homodimers, which needs to be further investigated.

It is interesting to note that cholesterol molecules bind to LYCHOS in the cytosolic side, which indicates the possibility for LYCHOS to sense concentrations of cholesterol in the cytosolic layer of the lysosome membrane. This is in consistence with the observation that OSBP, found at ER-lysosome membrane contact sites, transports cholesterol from the ER to the cytosolic layer of lysosome membrane and activates mTORC1^16^. By contrast, NPC1 suppresses mTORC1 by lowering the cholesterol levels. Cholesterol accumulates both in the lysosomal lumen and on the lysosomal membrane in cells lacking NPC1, the latter is mediated by ER-to-lysosome transport via OSBP, which leads to constitutively increased mTORC1 signalling. This makes sense that signal for cholesterol abundance comes from ER where it is synthesized.

In a previous study, researchers used photo-crosslinkable cholesterol analogs called LKM38 and KK231 to label recombinantly expressed LYCHOS. They discovered that both LKM38 and KK231 could be labeled onto TM1 of LYCHOS, specifically Glu48 and Cys55, respectively^19^. However, it is uncertain whether these cholesterol analogs bind in the same way as our structure suggests. This uncertainty arises because the side chains of Glu48 and Cys55 are positioned away from the cholesterol binding sites and are obstructed by another LYCHOS molecule (Extended Data Fig. 11a), preventing them from being labeled by LKM38 and KK231. Upon analyzing the structures of LKM38 and KK231, we observed that both have a bulky tail (Extended Data Fig. 11b-d). However, our structures suggest that cholesterol molecules bind in the cleft between PLD and GLD, with the hydrophobic tail penetrating deep into the pocket. This suggests that the bulky tails of LKM38 and KK231 may hinder their binding in this specific site. Consequently, it is not possible for LKM38 and KK231 to bind in the CBC. Nevertheless, apart from the canonical homodimer of LYCHOS, our structures also reveal another state: a non-canonical homodimer mediated by TM6 (Extended Data Fig. 9). In this state, TM1 is not buried by another LYCHOS molecule and is exposed in the membrane. Therefore, it is plausible that LKM38 and KK231 may interact with TM1 in some manner and label Glu48 and Cys55, respectively. In conclusion, caution should be taken when using photo-crosslinkable cholesterol analogs in future studies and LYCHOS shows multiple conformations.

Inhibition of mTORC1 with rapamycin is a promising therapy for age-related diseases. Many of the side effects of rapamycin and its analogs are now understood to be the result of off-target inhibition of a second mTOR complex, mTORC2. To address this issue, several laboratories and companies have started developing mTORC1-selective compounds with the aim of finding new ways to intervene in aging diseases and potentially extend lifespan^24^. Furthermore, it has been observed that mTORC1 activity is sustained by the accumulation of cholesterol on the lysosomal limiting membrane, supporting a process called senescence-associated secretory phenotype, which promotes aging^25^. Our studies on LYCHOS may contribute to the development of inhibitors that selectively target mTORC1 by blocking LYCHOS in its expanded state, which has therapeutic potential to treat aging diseases.

Recently, Ellisdon et al. reported structures of LYCHOS that differ significantly from our findings^23^. Notably, they did not observe the expanded states described in our manuscript, instead capturing only a closed state. Moreover, they were unable to obtain the structure of wild-type LYCHOS bound to cholesterol or its analogue CHS. Instead, they resorted to using an artificial mutant, LYCHOS(F352A/W678R), to achieve CHS binding. A critical methodological difference lies in the use of reducing agents. We incorporated these agents consistently throughout the purification process, whereas Ellisdon et al. added them only at the final stage. This distinction is crucial, as a large number of cysteine residues lies on the protein’s cytoplasmic face, contrasting with the absence of disulfide bonds in the lysosomal lumen. This distribution necessitates the continuous presence of reducing agents to maintain these cysteine residues in their native state and prevent denaturation. The consistent use of reducing agents throughout our purification process likely contributed to preserving LYCHOS in a more native-like conformation. This approach allowed us to capture a broader range of conformational states, including the expanded forms not observed by Ellisdon et al.

In summary, the structures of human LYCHOS were captured in the cholesterol bound contracted state, the lipids bound expanded state, and the TM6 mediated non-canonical expanded dimer state. Our studies reveal the overall structure and conformational plasticity of LYCHOS, uncover the cholesterol sensing mechanism by LYCHOS-mTORC1 pathway, lay the foundation for further mechanistic studies of LYCHOS and provide strategies for the design and development of LYCHOS-targeted therapeutics.

## Methods

### Cloning

UniProt (https://www.uniprot.org/) has access to the gene sequence used in this study. The full-length cDNA of GPR155 was cloned into pFastBac Dual, with the C-terminal mCherry, 6*His and twin-strep tags separated by a tobacco etch virus (TEV) protease cleavage site. A modified pET28a (+) with an N-terminal 6*His-MBP tag was used to clone the coding sequences of LED (538-658). Mutants of GPR155 and LED (538-658) were generated by site-directed mutagenesis.

### Cell Culture and Protein Expression

MBP-LED-WT was expressed in BL21(DE3). The bacteria were cultured in LB medium at 37℃ and supplemented with kanamycin (50 ug/ml), monitored until the Optical density at 600nm (OD600) reached 0.8-1.0. 0.1 mM IPTG was added and cultured overnight at 18 °C. After centrifugation, the bacteria were suspended using TBS (30 mM Tris, 3mM KCl, 140 mM NaCl,10% glycerol, pH7.4), then bacteria were flash-frozen in liquid nitrogen and stored at -80 °C until use.

LYCHOS proteins were expressed in SF9 cells using the baculovirus expression system. SF9 cells were cultured at 27 °C and virus was added when the density was 3× 10^6^ cells/ mL. Cells were collected after 60 h and washed by TBS, frozen with liquid nitrogen, and transferred to -80 °C for storage. Nprl2 and Nprl3 were also expressed in the SF9 expression system.

DEPDC5 was expressed in HEK293 cells, cultured at 37℃ and 5% CO2 with 1% fetal bovine serum (FBS). After reaching a cell density of 3×10^6^ cells/mL, 10% (v/v) virus was introduced and followed by the addition of 10mM sodium butyrate after 12 hours of infection. Cells were collected and re-suspended with TBS, then frozen with liquid nitrogen.

### Protein purification LYCHOS

The cells were thawed and resuspended in a buffer containing 50 mM HEPES at pH 7.4, 100 mM NaCl, 0.5 mM TCEP, and 1 mM phenylmethanesulfonylfluoride (PMSF). Following sonication using an ultrasonic cell crusher, the cells were solubilized for 3 hours with the addition of 1% LMNG/0.1% CHS. Subsequently, the lysate was centrifuged at 30,000 rpm for 40 minutes using a 45Ti rotor to remove insoluble material. The resulting supernatant was applied to Ni-IDA beads (Smart-Lifesciences) and washed with buffer A (50 mM HEPES at pH7.4, 100 mM NaCl, 0.5 mM TCEP, 0.005% LMNG/0.0005% CHS) containing appropriate concentrations of imidazole. After that, proteins were eluted with buffer A containing 300 mM imidazole. Dilute the eluent in equal volume buffer A, and applied to Strep-Tactin XT beads (IBA Lifesciences), then washed with buffer A. The resin was incubated overnight with TEV protease and eluted the following morning using buffer A; it was then concentrated and subjected to Superose6 increase column chromatography pre-equilibrated with buffer B (50 mM HEPES pH7.4, 100 mM NaCl, 0.5 mM TCEP, 0.002 %LMNG/0.0002 %CHS). Peak fractions containing LYCHOS were collected.

### GATOR1

Cells expressing Nprl2-Nprl3 were lysed in TBS containing 0.5 mM TCEP and 1 mM PMSF. After ultrasonic disruption, the lysate was centrifuged at 12,000 rpm for 30 minutes, and the supernatant was loaded onto Ni-IDA beads (Smart-Lifesciences). The beads were washed with buffer D (20 mM Tris, pH 8.0, 500 mM NaCl, 1% glycerol) containing 0.5 mM TCEP and appropriate concentrations of imidazole. Proteins were then eluted with buffer D containing 300 mM imidazole and 1 mM TCEP. The eluate was incubated with TEV protease overnight, concentrated, and then run on a Superose 6 increase column pre-equilibrated with buffer E (20 mM Tris, 150 mM NaCl, 1% glycerol, pH 8.0) plus 0.5 mM TCEP. The purification procedures for DEPDC5 with tag were identical, except without the use of TEV protease.

### MBP-LED and MBP-LED-Y551A

Each step is identical to that of DEPDC5, except that TCEP is used at double the concentration.

### Microscale Thermophoresis (MST) Assay

The GATOR1 binding affinities of LYCHOS and MBP-LED were quantified using MST. Purified GATOR1 proteins (10 μM) were exchanged into a labeling buffer and labeled with the dye NHS. The labeled protein was displaced by MST buffer containing 50 mM HEPES pH 7.4, 100 mM NaCl, and 0.05% (v/v) Tween-20. 16 gradient dilution concentrations of LYCHOS or MBP-LED were prepared. 10 ul labeled GATOR1 were mixed with 10 ul LYCHOS or MBP-LED. The samples in tubes 1 to 16 were loaded into premium capillary tube. Binding reactions were measured using a microscale thermophoresis instrument (Nano Temper Technologies GMBH) at 25℃, 40% MST power and 20% LED power. The K_D_ Fit function of the Nano Temper Analysis Software MO Affinity Analysis (V2.3) was used to fit the curve and calculate the value of the dissociation constant (Kd).

### Cryo-EM sample preparation and data acquisition

The protein samples were loaded onto quantifoil R 1.2/1.3 300 Holey Carbon films Au 300 mesh grids and blotted with filter paper for 3 s under 100% humidity at 6 °C before being plunged into liquid ethane with a FEI Vitrobot Mark IV. Data collection was accomplished using a Falcon4 direct electron detection camera and a 300 kV Titan Krios G4 electron microscopy. The movies were captured using a calibrated magnification of 96k x with a pixel size of 0.404 Å. Total dose of 49.9 e/Å^2^ was used.

### Cryo-EM image analysis

The detailed image processing workflows are illustrated in Extended Data Figs. 2c, 3c, 7c, 8c, and 9c. The movies were twofold binned to a pixel size of 0.808 Å in motion-correction using MotionCor2^26^. cryoSPARC^27^ was employed for all subsequent classification and reconstruction. Contrast transfer function (CTF) parameters were estimated by Patch CTF estimation. Templates were used for template picking. After several iterations of 2D Classification, particles from good classes were used for 3D ab initio reconstruction to generate volumes, which were used as input for heterogeneous refinement or non-uniform refinement afterwards. For LYCHOS_WT_ and LYCHOS_Y57A_ datasets, particles were further classified using three-dimensional (3D) classification, and good classes were refined using NU refinement.

### Model building and refinement

AlphaFold2 was used to predict the initial model of LYCHOS. Subsequently, the initial model was fitted into the cryo-EM map using UCSF Chimera^28^ and manually rebuilt utilizing Coot^29^. The model was further refined using phenix.real_space_refine in Phenix^30^. Images were generated using PyMOL (Schrödinger, LLC), UCSF Chimera^28^, and ChimeraX^31^.

### Molecular dynamics (MD) simulation analysis

The CHARMM-GUI^32^ website is used to process the protein file. The protein complexes were inserted in a lipid bilayer composed of palmitoyl-oleoyl-phosphatidyl-choline (POPC), palmitoyl-oleoyl-phosphatidyl-ethanolamine (POPE), palmitoyl-oleoyl-phosphatidyl-inositol (POPI), sphingomyelin (SM), cholesterol and lysobisphosphatidic acid (LBPA, BMP) with a molar ratio of 30:11:7:15:30:7, similar to lysosome membrane components previously reported^33^. The system was then solvated with the TIP3 water model, and 150 mM KCl was added to mimic physiological conditions. A periodic rectangular box with approximate dimensions of 13.3 nm x 13.3 nm x 12.9 nm was applied, which contains around 200,000 atoms. The CHARMM36m^34^ force field was used to describe proteins and lipids. Prepared systems were first minimized using 5000 steps of a steepest descent algorithm. Next, 100 ps was used to equilibrate the system at 310 K using multi-step isothermal–isovolumetric (NVT) and isothermal–isobaric (NPT) conditions while decreasing the restraints at each step, and a 1000 ns MD simulation was conducted with 2 fs time integration steps at a constant temperature of 310 K using the Gromacs 2023^35^ software package. Analyses were performed using the Gromacs 2023 package and the visual molecular dynamics (VMD) program.

### GATOR1 recruitment assay

HEK293 cells were transiently co-transfected with LYCHOS-Rluc and GATOR1 heterotrimers, including NPRL2-YFP, NPRL3, and DEPDC5. Twenty-four hours following transfection, the cells were distributed into 96-well microtitre plates at a density of 5 x 10⁴ cells per well. Following a 24-hour incubation period at 37°C, the cells were washed twice with Tyrode’s buffer and subsequently stimulated with the indicated concentrations of ligands. Prior to recording light emission, the luciferase substrate coelenterazine-h (5 µM) was added. This was done using a Mithras LB940 microplate reader (Berthold Technologies) equipped with a BRET filter set. The BRET signal was calculated as the luminescence ratio at 530 nm/485 nm. The change in BRET signal due to ligand addition was recorded as ΔBRET.

### Cell lysis and immunoblotting

For HEK293T cells, 0.3 million cells/well were seeded on a 6 well plate. The following day, cells were rinsed with DMEM added 8% FBS. The FLAG-S6K1 and corresponding LYCHOS-mCherry plasmid were transfected to cells with PEI. After 48 h, cells were stimulated with 100 µM cholesterol in complex with 0.1% MCD for 30 min. Then cells were washed once with PBS and lysed in buffer (1% NP40, 3 mM PMSF, 1× phosphatase inhibitor cocktail, and 1× EDTA-free protease inhibitor cocktail) for 30 min. Cell lysates were cleared by centrifugation in a microcentrifuge at 12000 rpm for 10 min at 4 °C. Cell lysate samples were measured by BCA and samples of equalized concentration were prepared for SDS-PAGE by addition of 5x sample loading buffer. Indicated proteins were detected by immunoblotting.

### Data availability

The atomic model coordinates and cryo-EM maps have been deposited in the Protein Data Bank (PDB) and the Electron Microscopy Data Bank (EMDB), respectively, under the accession codes: 9JBE and EMD-61311 (LYCHOS_WT_C_), 9JBG and EMD-61313 (LYCHOS_WT_E_), 9JBF and EMD-61312 (LYCHOS_Y57A_C_), 9JBI and EMD-61315 (LYCHOS_Y57A_TM6_E_), 9JBJ and EMD-61316 (LYCHOS_Y57A/R61A_E_) and 9JBH and EMD-61314 (LYCHOS_PLD_). Any additional information required to reanalyze the data reported in this paper is available from the Lead Contact upon request.

## Acknowledgements

We thank the cryo-EM facility from Shuimu BioSciences for data collection. This work is also supported by the Cryo-EM platform of Peking University Health Science Center with the assistance of Dr. Dandan Chen. This work was supported by grants including the National Natural Science Foundation of China (grant 32171224 to L.L.), the Fundamental Research Funds for the Central Universities (BMU2022RCZX008 to L.L.) and Lam Chung Nin Foundation for Systems Biomedicine.

## Author contributions

L.L. and J.S. conceived the project. S.Y. made the constructs, purified proteins, and performed biochemical experiments. J.D., W.W. and J.W. performed cell assays. Z.P. prepared cryo-EM grids. A.Y. and Z.D. assisted in the protein purification. L.L. processed the cryo-EM data, built models and performed modelling and refinement. L.L. performed and analyzed the molecular dynamics simulation. Y.Y. provided helpful discussion. L.L. wrote the manuscript with input from all authors.

---

**Extended Data Fig. 1.**
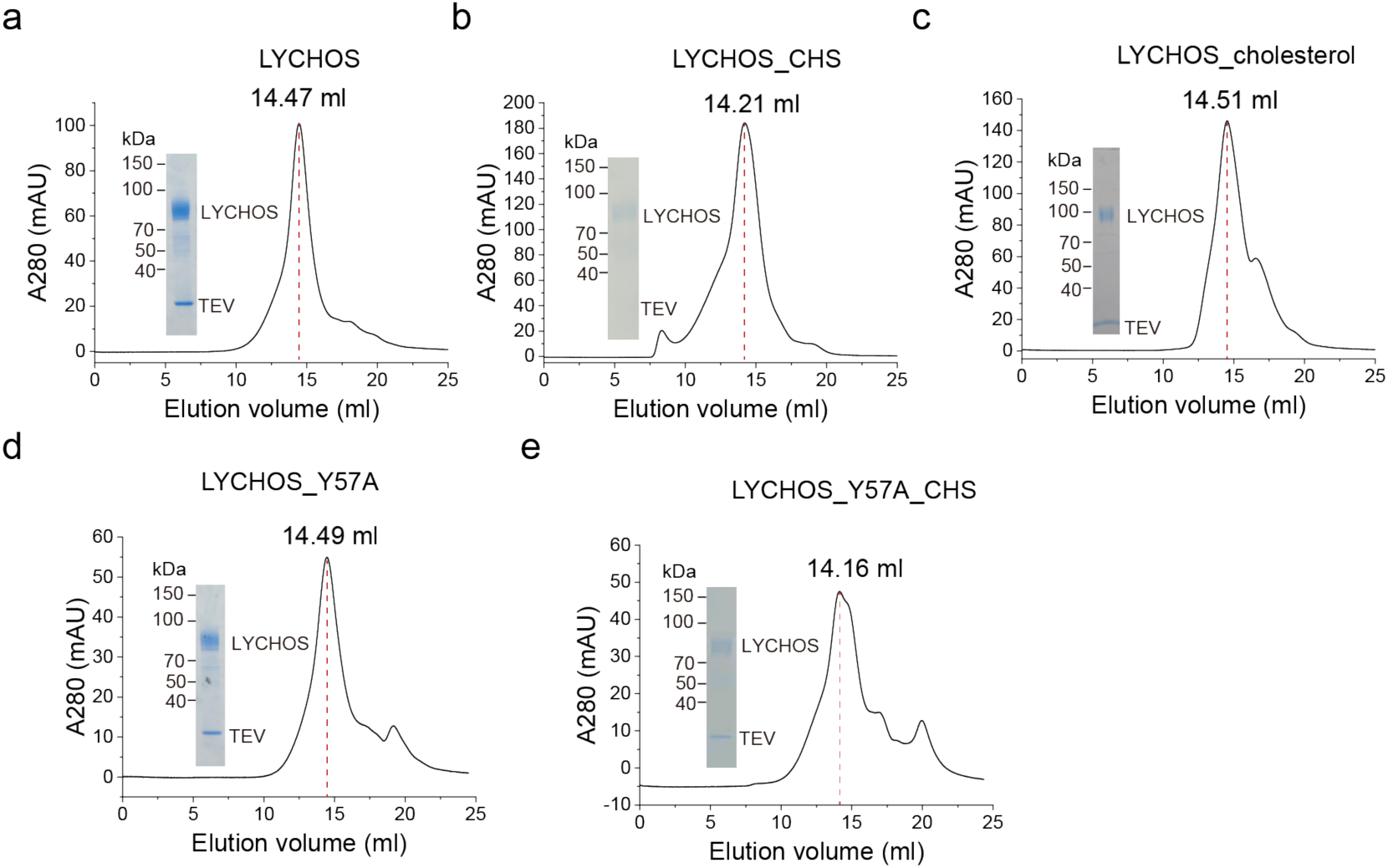
| Purification of LYCHOS. a-g, Size-exclusion chromatography on Superose 6 and Coomassie-blue-staining SDS-PAGE results of LYCHOS, LYCHOS with CHS, LYCHOS with cholesterol, LYCHOS Y57A mutant and LYCHOS Y57A mutant with CHS, respectively. n=2 or 3 independent experiments.

**Extended Data Fig. 2.**
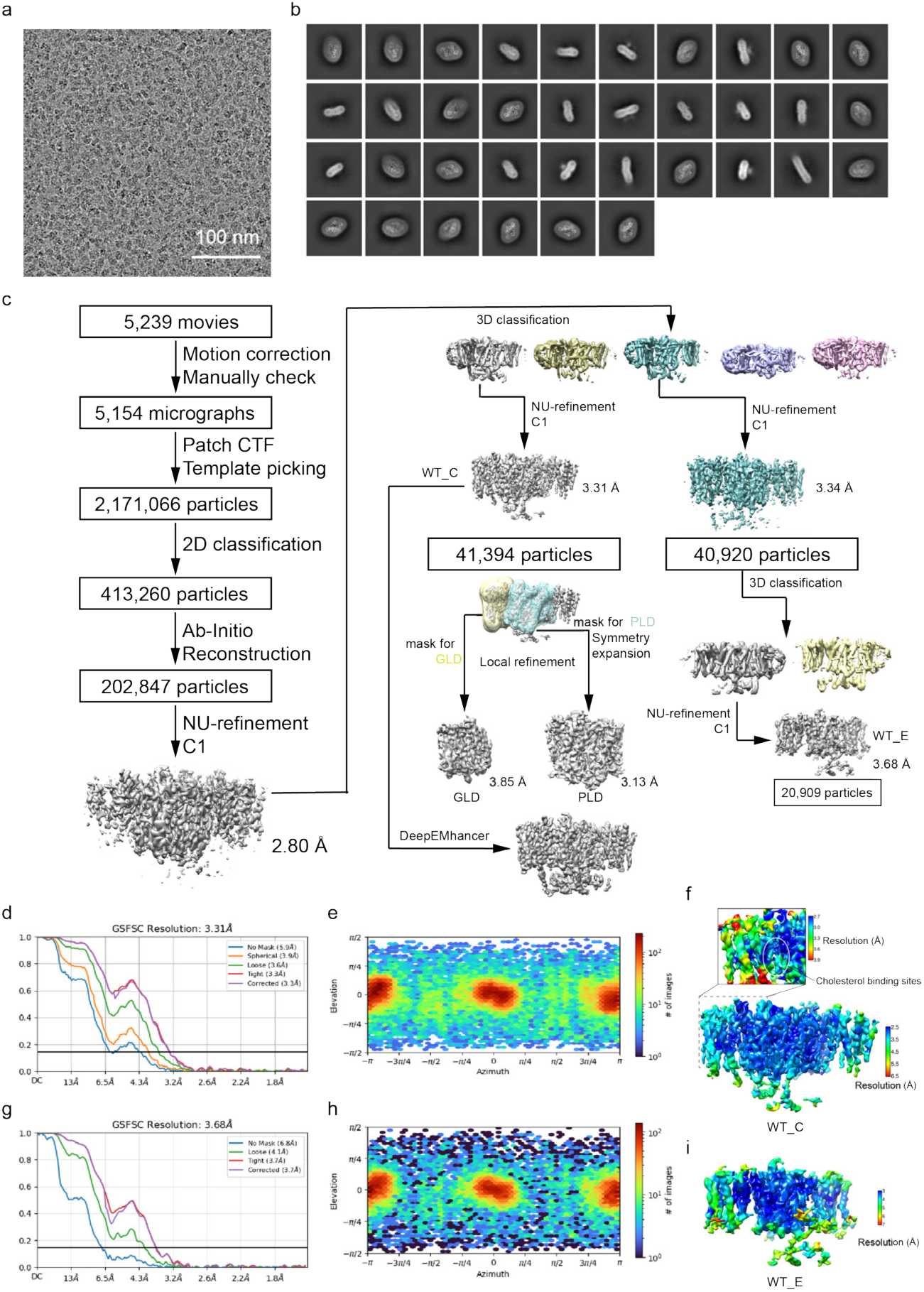
| Cryo-EM image analysis of LYCHOS_WT_. a, Representative raw micrograph (5,239 in total) of LYCHOS_WT_ in the presence of CHS. Scale bar, 100 nm. b, 2D-class averages of LYCHOS_WT_ in the presence of CHS. c, Cryo-EM data processing workflow of LYCHOS_WT_ in the presence of CHS. d, Gold-standard Fourier Shell Correlation (FSC) of the refined map of LYCHOS_WT_C_. e, Angular distribution of the final reconstruction of LYCHOS_WT_C_. f, A view of the local resolution map of the LYCHOS_WT_C_. Scale bar, 2.5-6.5 Å. The local resolution map of the cholesterol binding sites is shown as an insert. Scale bar, 2.7-3.9 Å. The local resolution around cholesterol indicated by a white circle is higher than 3.3 Å. g, Gold-standard Fourier Shell Correlation (FSC) of the refined map of LYCHOS_WT_E_. h, Angular distribution of the final reconstruction of LYCHOS_WT_E_. i, A view of the local resolution map of the LYCHOS_WT_E_. Scale bar, 3-7 Å.

**Extended Data Fig. 3.**
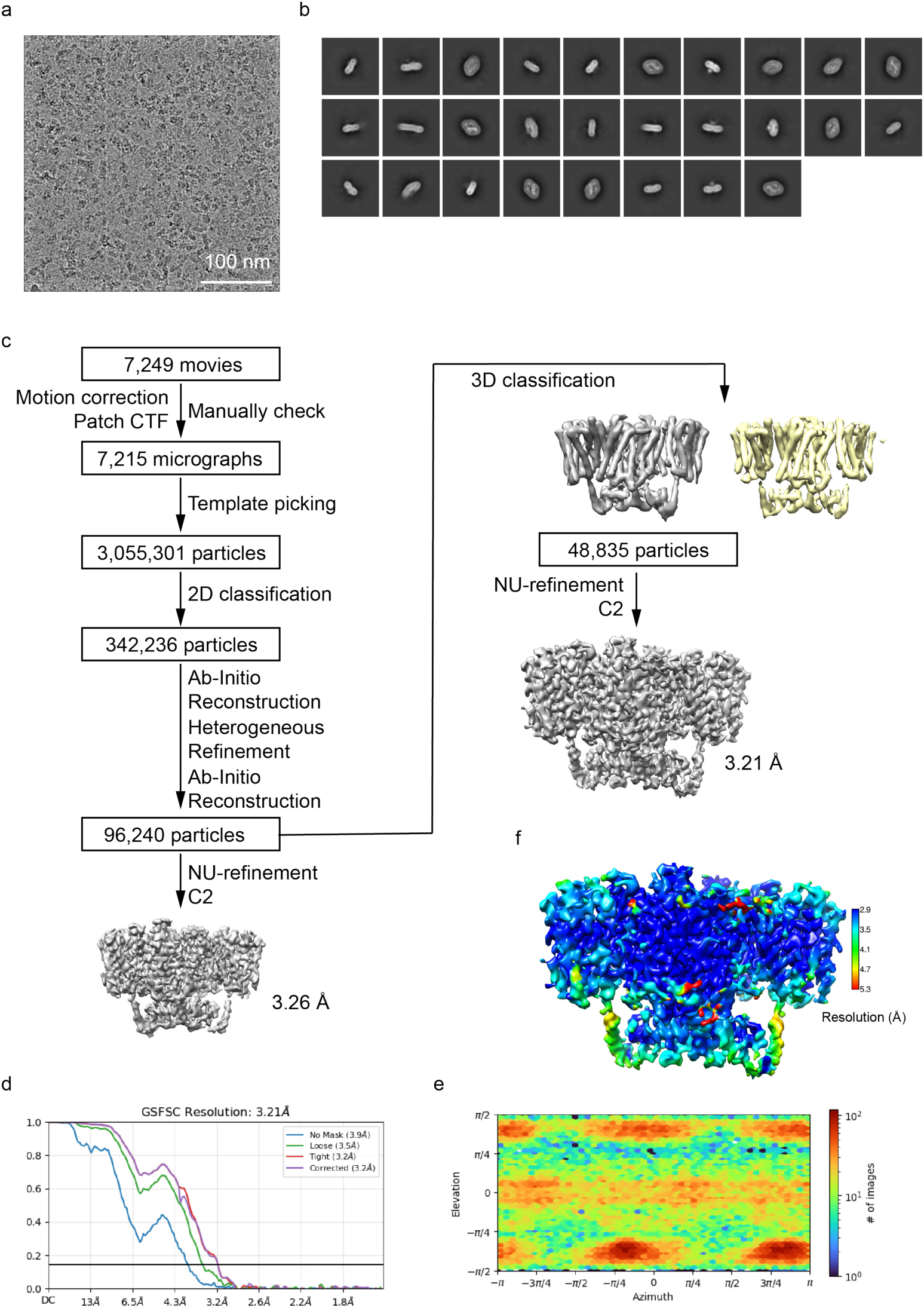
| Cryo-EM image analysis of LYCHOS_Y57A_C_. a, Representative raw micrograph (7,249 in total) of LYCHOS_Y57A_ in the presence of CHS. Scale bar, 100 nm. b, 2D-class averages of LYCHOS_Y57A_ in the presence of CHS. c, Cryo-EM data processing workflow of LYCHOS_Y57A_ in the presence of CHS. d, Gold-standard Fourier Shell Correlation (FSC) of the refined map shown in c. e, Angular distribution of the final reconstruction. f, A view of the local resolution map of the LYCHOS_Y57A_C_. Scale bar, 2.9-5.3 Å.

**Extended Data Fig. 4.**
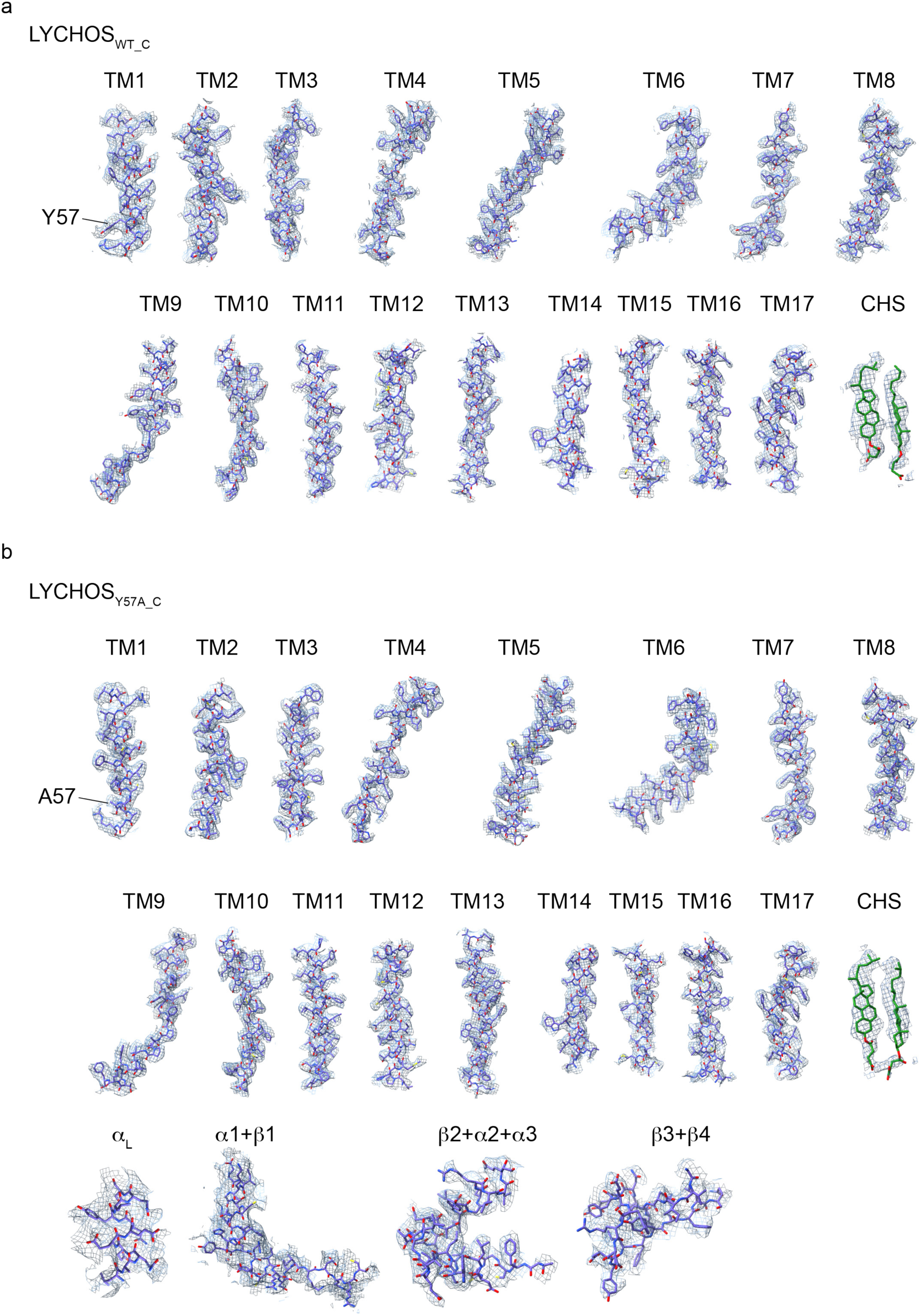
| Representative cryo-EM density maps. a, Cryo-EM density maps and models of transmembrane helices and CHS molecules are shown in the LYCHOS_WT_C_ using its global map with map threshold at 0.15. b, Cryo-EM density maps and models of transmembrane helices, DEP and CHS molecules are shown in the LYCHOS_Y57A_C_ with map threshold at 0.15, 0.10 and 0.22, respectively.

**Extended Data Fig. 5.**
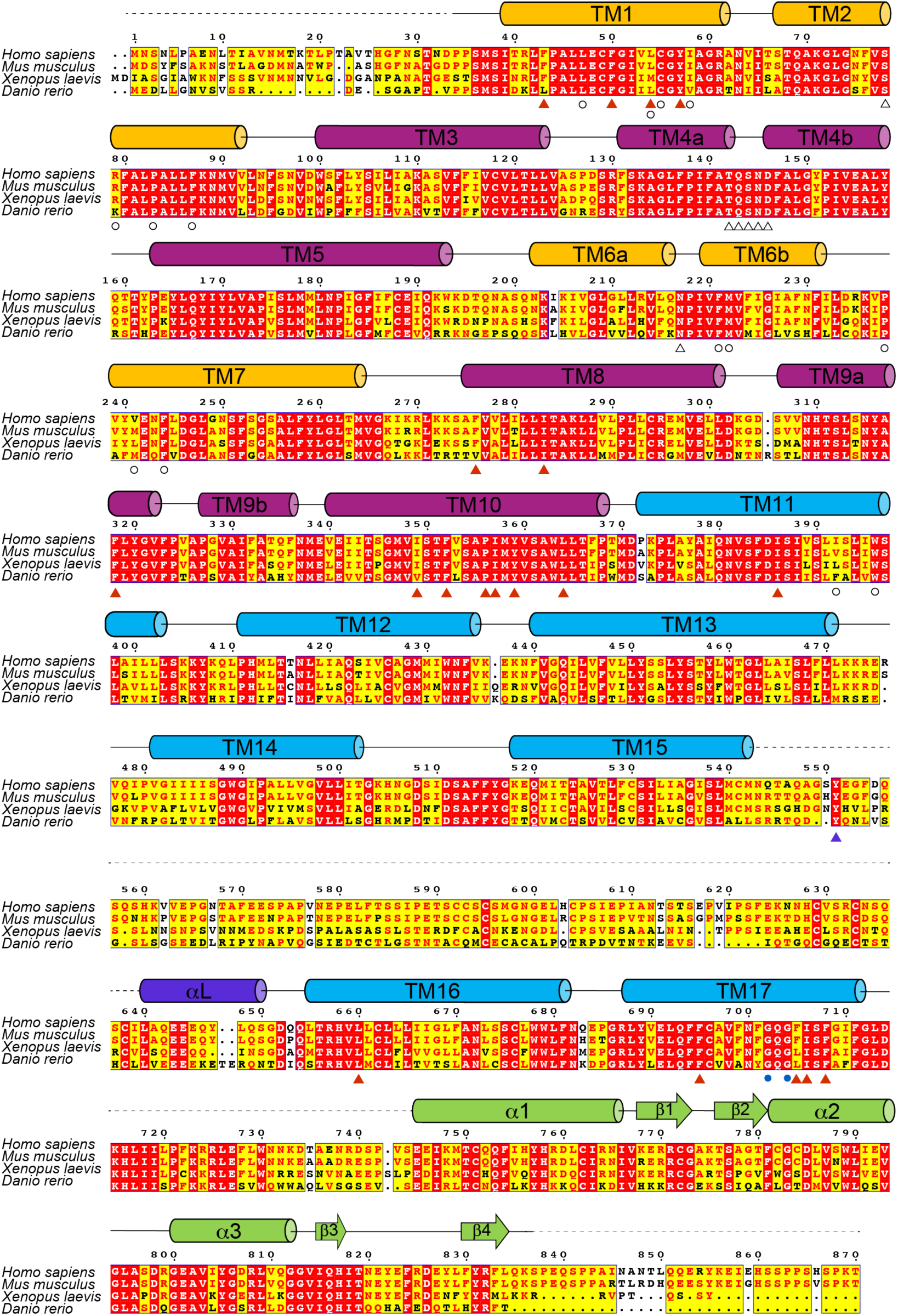
| Multiple sequence alignment of LYCHOS orthologues. Elements of secondary structure are shown above the sequence with the same color as in Fig. 1a. Highly conserved residues are shaded in red. Relatively conserved residues are shaded in yellow and colored in red. Residues for binding with the cholesterol molecules are denoted by red triangle. Residues for bending of TM17 are denoted by blue circle. Residues for binding with lipids are denoted by hollow circle. Proposed substrate binding residues are denoted by hollow triangle. Residues for binding with GATOR1 are denoted by blue triangle.

**Extended Data Fig. 6.**
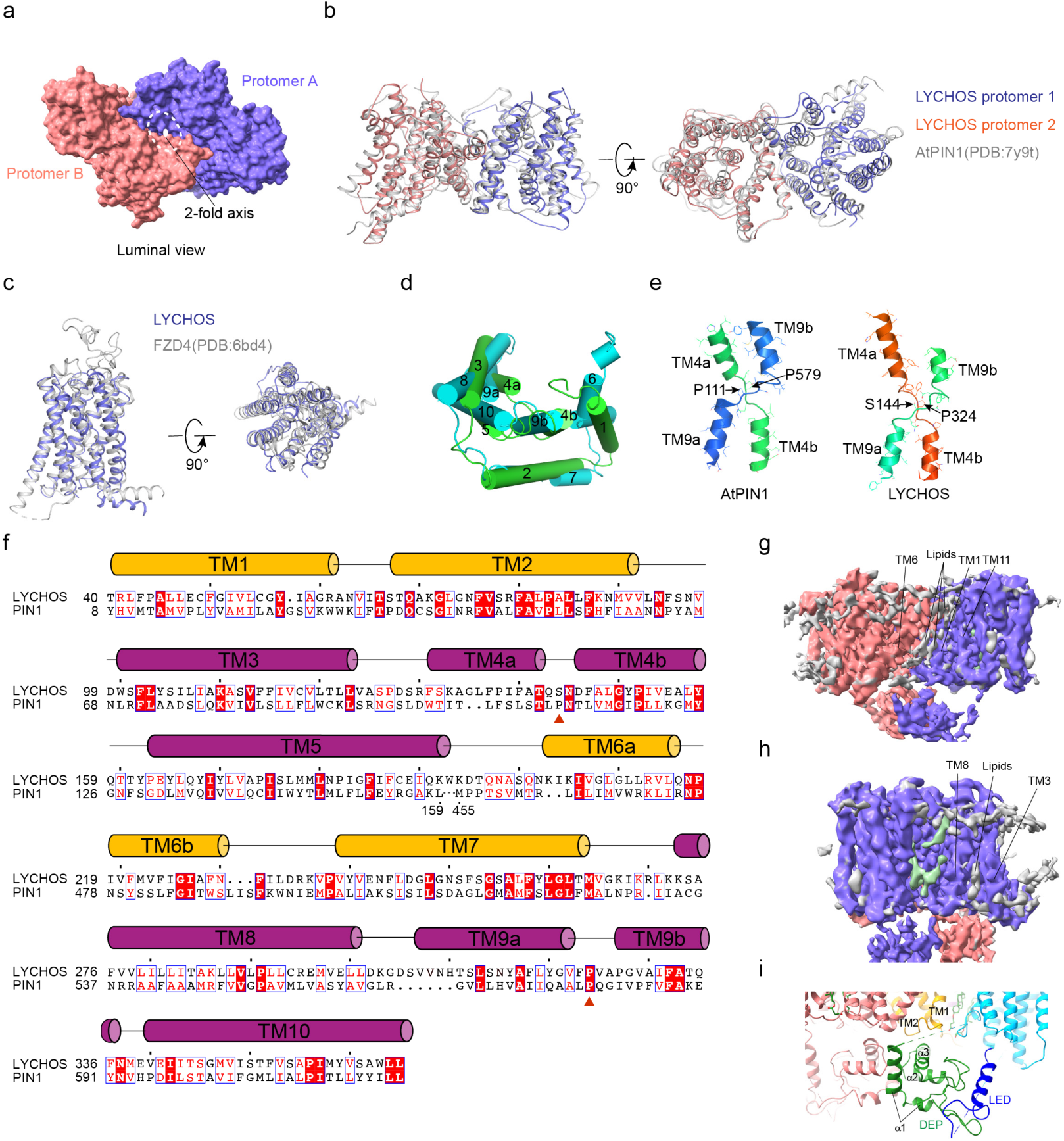
| Structure comparisons of LYCHOS, PIN1 and FZD4 and lipids binding LYCHOS. a, The luminal-facing cavity at the dimer interface shown in luminal view. The dashed circle indicates the bowl-shaped luminal-facing cavity. b, Superimposition of LYCHOS PLD and PIN1 structures in different orientations. c, Superimposition of LYCHOS GLD and FZD4 structures in different orientations. d, The first and last five TMs of LYCHOS are structurally inverted repeats. TM6-10 merges with TM1-5. e, TM4 and TM9 of LYCHOS form a crossover structure as PIN1. Side chains are shown as lines. Conserved residues for bending are indicated with an arrow. f, Sequence alignment of LYCHOS and PIN1, secondary structures of LYCHOS are shown above the sequence with the same color as in Fig. 1b. Conserved residues for bending are denoted by red triangle. g, Position of lipids in the groove between the two protomers. h, The lipid between TM3 and TM8. i, The dimer interface of DEP and its interactions with LED and TMs.

**Extended Data Fig. 7.**
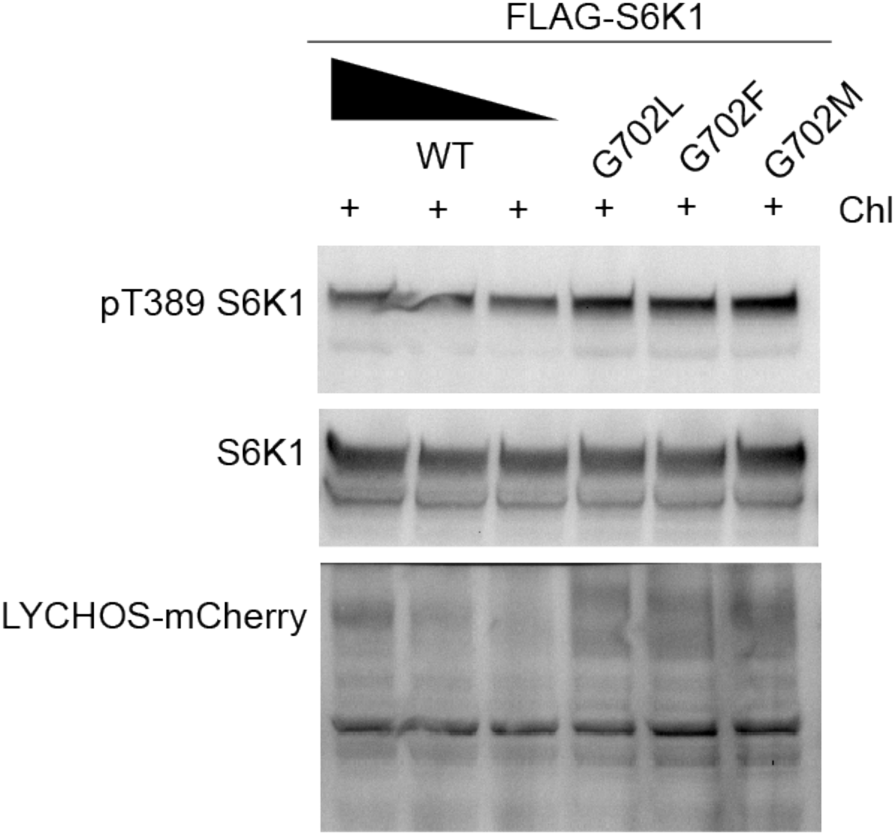
| Bulky side chain mutations of LYCHOS Gly702 result in elevated activation of mTORC1. HEK293T were transfected with FLAG-S6K1 along with decreasing amount of mCherry-tagged LYCHOS WT and G702 mutants. Cells were stimulated with 100 µM cholesterol (Chl) in complex with 0.1% MCD, followed by immunoblotting for the phosphorylation state and levels of the indicated proteins.

**Extended Data Fig. 8.**
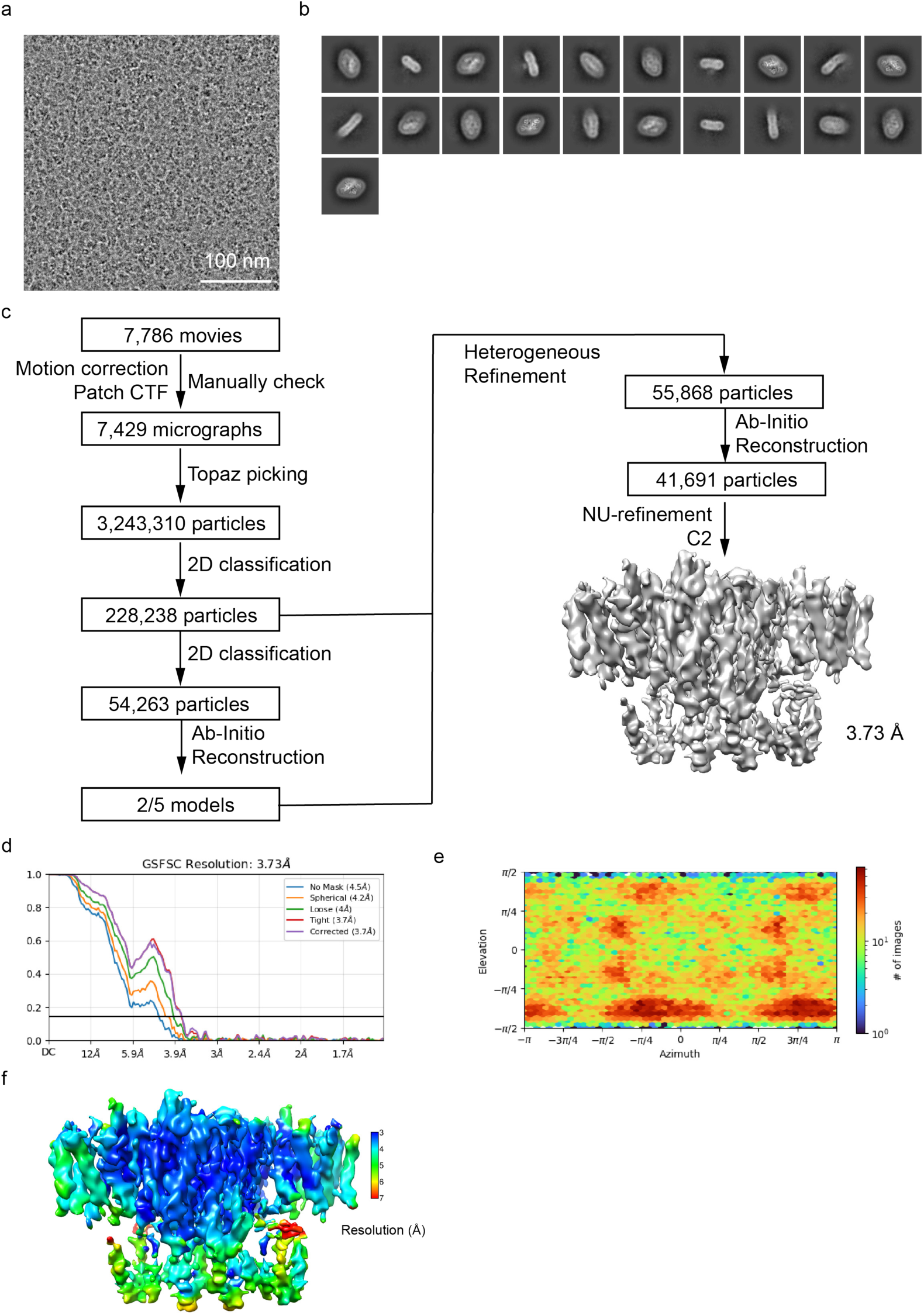
| Cryo-EM image analysis of LYCHOS_Y57A/R61A_. a, Representative raw micrograph (7,786 in total) of LYCHOS_Y57A/R61A_ in the presence of CHS. Scale bar, 100 nm. b, 2D-class averages of LYCHOS_Y57A/R61A_ in the presence of CHS. c, Cryo-EM data processing workflow of LYCHOS_Y57A/R61A_ in the presence of CHS. d, Gold-standard Fourier Shell Correlation (FSC) of the refined map shown in c. e, Angular distribution of the final reconstruction. f, A view of the local resolution map of the LYCHOS_Y57A/R61A_. Scale bar, 3-7 Å.

**Extended Data Fig. 9.**
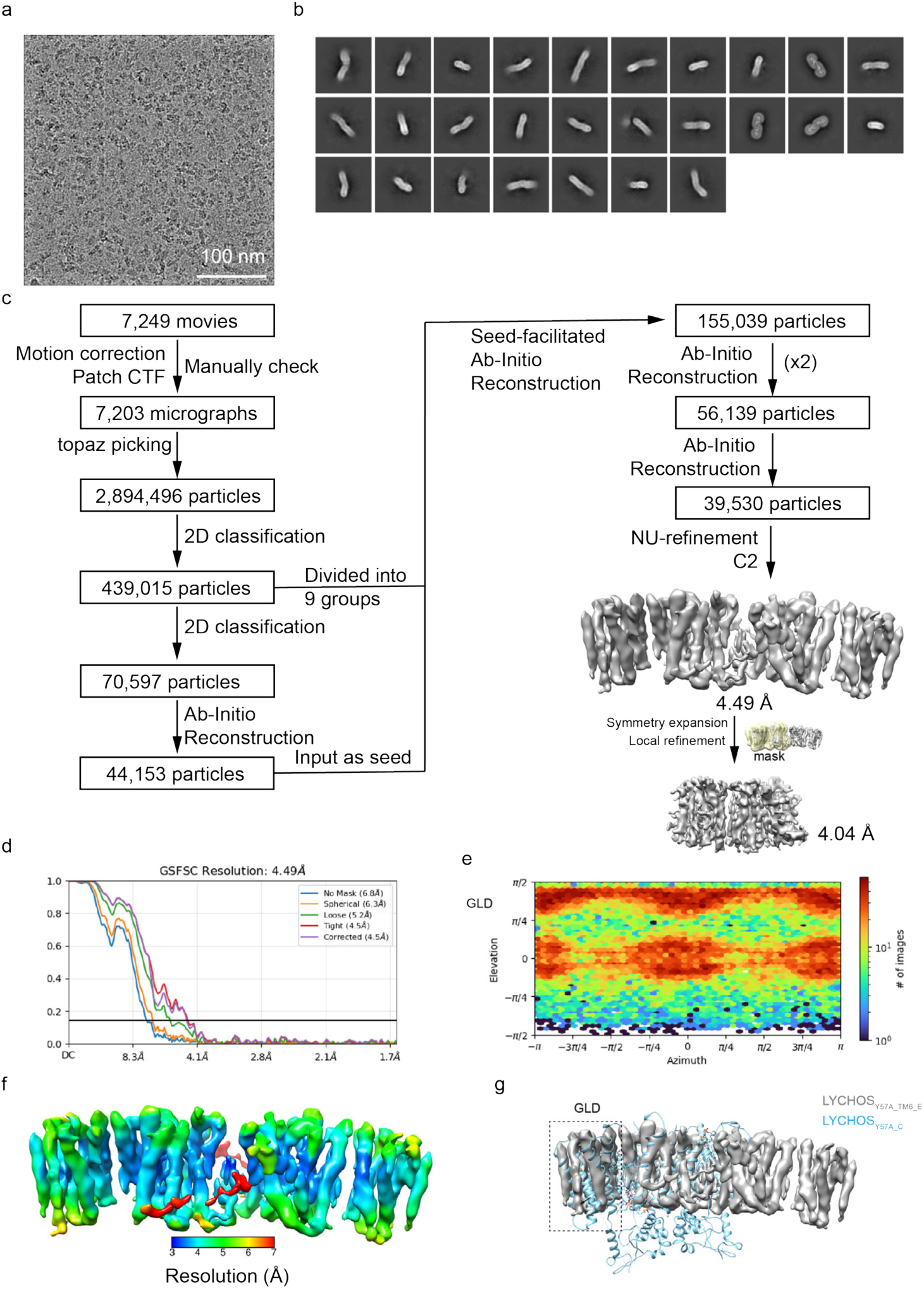
| Cryo-EM image analysis of LYCHOS_Y57A_TM6_. a, Representative raw micrograph (7,249 in total) of LYCHOS_Y57A_ in the presence of CHS. Scale bar, 100 nm. b, 2D-class averages of LYCHOS_Y57A_TM6_ in the presence of CHS. c, Cryo-EM data processing workflow of LYCHOS_Y57A_TM6_ in the presence of CHS. d, Gold-standard Fourier Shell Correlation (FSC) of the refined map shown in c. e, Angular distribution of the final reconstruction. f, A view of the local resolution map of the LYCHOS_Y57A_TM6_. Scale bar, 3-7 Å. g, Superimposition of the LYCHOS_Y57A_C_ structure with the map of LYCHOS_Y57A_TM6_E_. The solid arrow indicates the translation of GLD.

**Extended Data Fig. 10.**
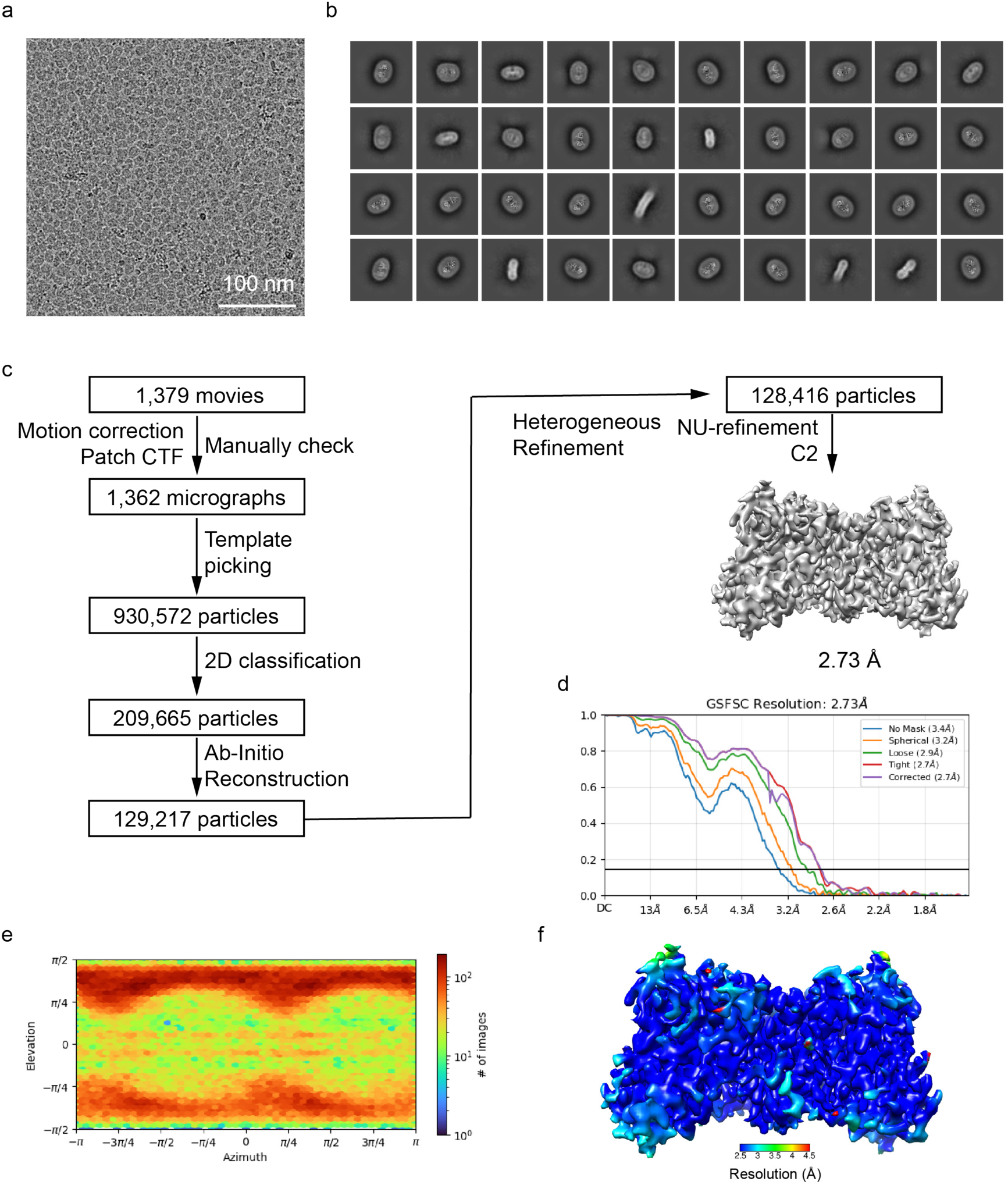
| Cryo-EM image analysis of LYCHOS_PLD_. a, Representative raw micrograph (1,379 in total) of LYCHOS_PLD_ in the presence of CHS. Scale bar, 100 nm. b, 2D-class averages of LYCHOS_PLD_ in the presence of CHS. c, Cryo-EM data processing workflow of LYCHOS_PLD_ in the presence of CHS. d, Gold-standard Fourier Shell Correlation (FSC) of the refined map shown in c. e, Angular distribution of the final reconstruction. f, A view of the local resolution map of the LYCHOS_PLD_. Scale bar, 2.5-4.5 Å.

**Extended Data Fig. 11.**
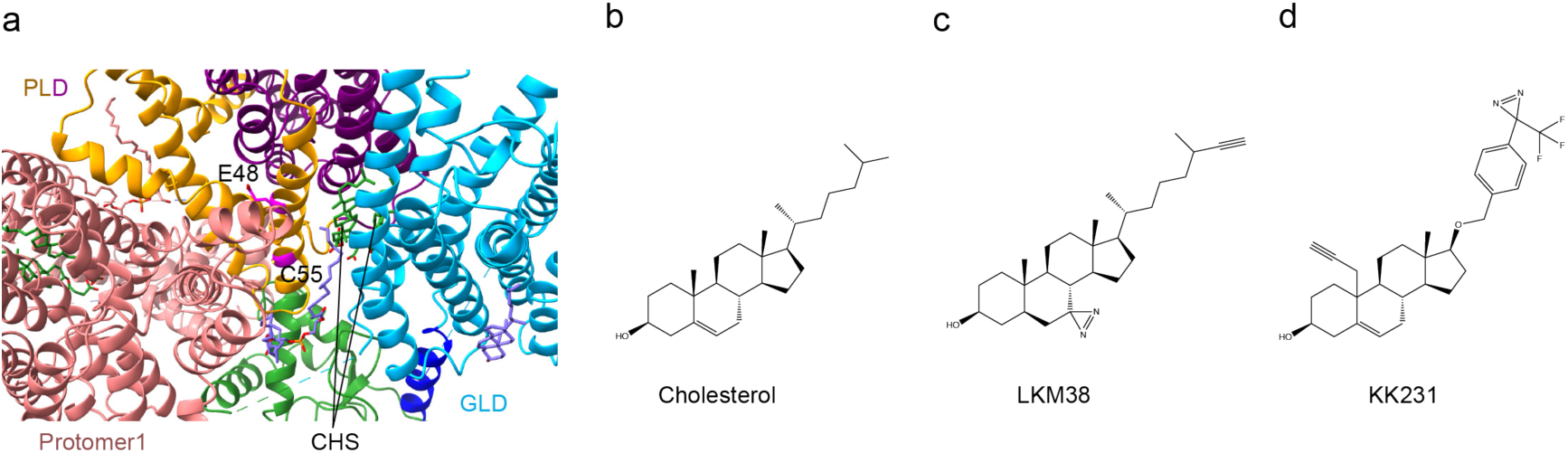
| Canonical dimer of LYCHOS hinders the labeling of Glu48 and Cys55 by cholesterol analog probes. a, Structural model of LYCHOS_WT_C_ shown in cartoon representation. Domains are colored as in Fig. 1f. Glu48, Cys55 and CHS molecules are shown in sticks. b-d, 2D structures of cholesterol, LKM38 and KK231. The hydrophobic tails of LKM38 and KK231 are larger than cholesterol.

**Extended Data Table 1.**
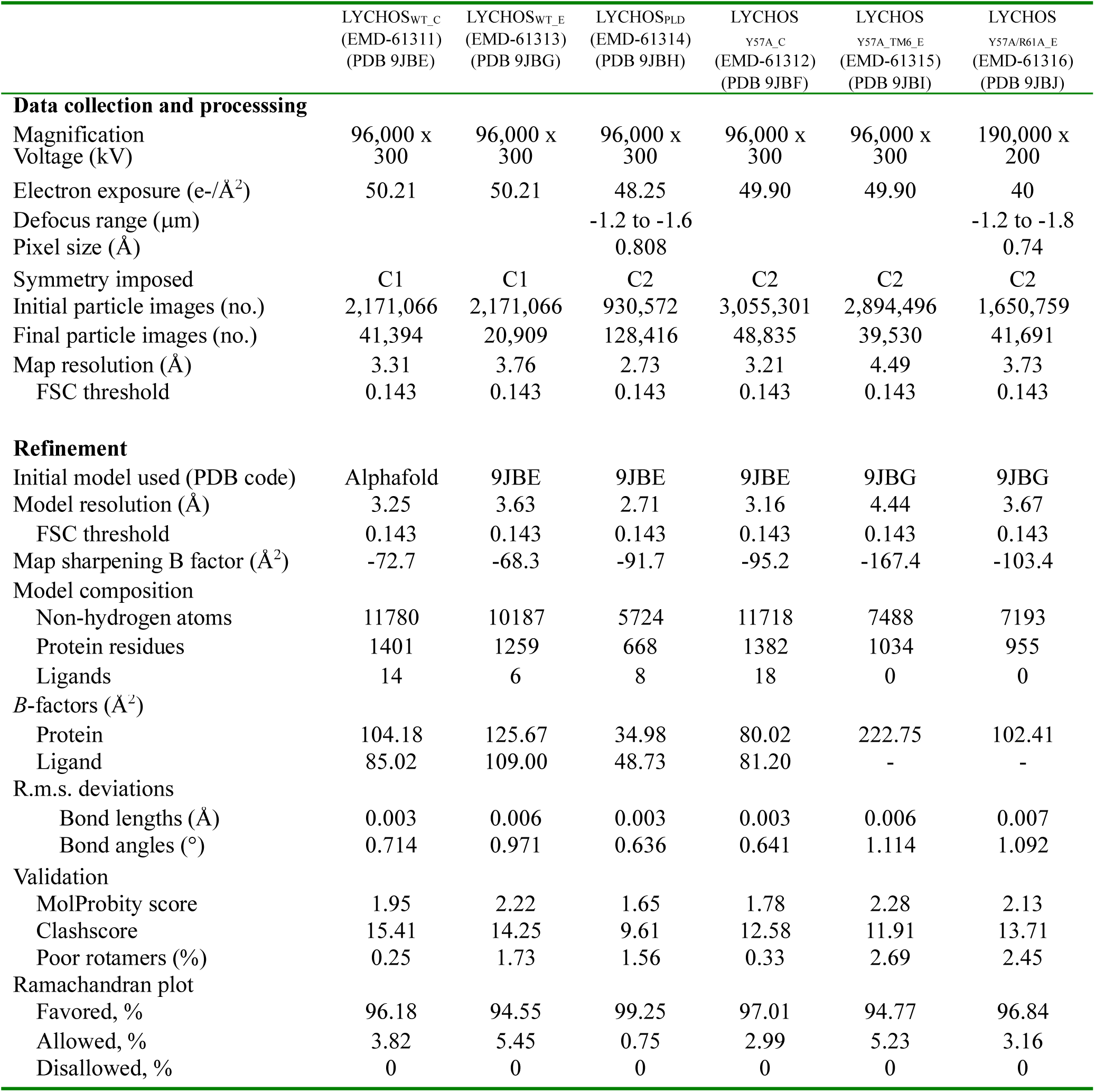
| Cryo-EM data collection, refinement and validation statistics

## References

1 Kim, E., Goraksha-Hicks, P., Li, L., Neufeld, T. P. & Guan, K. L. Regulation of TORC1 by Rag GTPases in nutrient response. Nat Cell Biol 10, 935–945, doi:10.1038/ncb1753 (2008).

2 Sancak, Y. et al. The Rag GTPases bind raptor and mediate amino acid signaling to mTORC1. Science 320, 1496–1501, doi:10.1126/science.1157535 (2008).

3 Perera, R. M. & Zoncu, R. The Lysosome as a Regulatory Hub. Annu Rev Cell Dev Biol 32, 223–253, doi:10.1146/annurev-cellbio-111315-125125 (2016).

4 Sancak, Y. et al. Ragulator-Rag complex targets mTORC1 to the lysosomal surface and is necessary for its activation by amino acids. Cell 141, 290–303, doi:10.1016/j.cell.2010.02.024 (2010).

5 Anandapadamanaban, M. et al. Architecture of human Rag GTPase heterodimers and their complex with mTORC1. Science 366, 203–210, doi:10.1126/science.aax3939 (2019).

6 Yang, H. et al. Mechanisms of mTORC1 activation by RHEB and inhibition by PRAS40. Nature 552, 368–373, doi:10.1038/nature25023 (2017).

7 Bar-Peled, L. et al. A Tumor suppressor complex with GAP activity for the Rag GTPases that signal amino acid sufficiency to mTORC1. Science 340, 1100–1106, doi:10.1126/science.1232044 (2013).

8 Tsun, Z. Y. et al. The folliculin tumor suppressor is a GAP for the RagC/D GTPases that signal amino acid levels to mTORC1. Mol Cell 52, 495–505, doi:10.1016/j.molcel.2013.09.016 (2013).

9 Lawrence, R. E. et al. Structural mechanism of a Rag GTPase activation checkpoint by the lysosomal folliculin complex. Science 366, 971–977, doi:10.1126/science.aax0364 (2019).

10 Shen, K. et al. Architecture of the human GATOR1 and GATOR1-Rag GTPases complexes. Nature 556, 64–69, doi:10.1038/nature26158 (2018).

11 Egri, S. B. et al. Cryo-EM structures of the human GATOR1-Rag-Ragulator complex reveal a spatial-constraint regulated GAP mechanism. Mol Cell 82, 1836–1849 e1835, doi:10.1016/j.molcel.2022.03.002 (2022).

12 Wolfson, R. L. et al. KICSTOR recruits GATOR1 to the lysosome and is necessary for nutrients to regulate mTORC1. Nature 543, 438–442, doi:10.1038/nature21423 (2017).

13 Luo, J., Yang, H. & Song, B. L. Mechanisms and regulation of cholesterol homeostasis. Nat Rev Mol Cell Biol 21, 225–245, doi:10.1038/s41580-019-0190-7 (2020).

14 Meng, Y., Heybrock, S., Neculai, D. & Saftig, P. Cholesterol Handling in Lysosomes and Beyond. Trends Cell Biol 30, 452–466, doi:10.1016/j.tcb.2020.02.007 (2020).

15 Castellano, B. M. et al. Lysosomal cholesterol activates mTORC1 via an SLC38A9-Niemann-Pick C1 signaling complex. Science 355, 1306–1311, doi:10.1126/science.aag1417 (2017).

16 Lim, C. Y. et al. ER-lysosome contacts enable cholesterol sensing by mTORC1 and drive aberrant growth signalling in Niemann-Pick type C. Nat Cell Biol 21, 1206–1218, doi:10.1038/s41556-019-0391-5 (2019).

17 Rebsamen, M. et al. SLC38A9 is a component of the lysosomal amino acid sensing machinery that controls mTORC1. Nature 519, 477–481, doi:10.1038/nature14107 (2015).

18 Wang, S. et al. Metabolism. Lysosomal amino acid transporter SLC38A9 signals arginine sufficiency to mTORC1. Science 347, 188–194, doi:10.1126/science.1257132 (2015).

19 Shin, H. R. et al. Lysosomal GPCR-like protein LYCHOS signals cholesterol sufficiency to mTORC1. Science 377, 1290–1298, doi:10.1126/science.abg6621 (2022).

20 Su, N. et al. Structures and mechanisms of the Arabidopsis auxin transporter PIN3. Nature 609, 616–621, doi:10.1038/s41586-022-05142-w (2022).

21 Ung, K. L. et al. Structures and mechanism of the plant PIN-FORMED auxin transporter. Nature 609, 605–610, doi:10.1038/s41586-022-04883-y (2022).

22 Yang, Z. et al. Structural insights into auxin recognition and efflux by Arabidopsis PIN1. Nature 609, 611–615, doi:10.1038/s41586-022-05143-9 (2022).

23 Bayly-Jones, C. et al. LYCHOS is a human hybrid of a plant-like PIN transporter and a GPCR. Nature, doi:10.1038/s41586-024-08012-9 (2024).

24 Dumas, S. N. & Lamming, D. W. Next Generation Strategies for Geroprotection via mTORC1 Inhibition. J Gerontol A Biol Sci Med Sci 75, 14–23, doi:10.1093/gerona/glz056 (2020).

25 Roh, K. et al. Lysosomal control of senescence and inflammation through cholesterol partitioning. Nat Metab 5, 398–413, doi:10.1038/s42255-023-00747-5 (2023).

26 Zheng, S. Q. et al. MotionCor2: anisotropic correction of beam-induced motion for improved cryo-electron microscopy. Nature methods 14, 331–332, doi:10.1038/nmeth.4193 (2017).

27 Punjani, A., Rubinstein, J. L., Fleet, D. J. & Brubaker, M. A. cryoSPARC: algorithms for rapid unsupervised cryo-EM structure determination. Nature methods 14, 290–296, doi:10.1038/nmeth.4169 (2017).

28 Pettersen, E. F. et al. UCSF Chimera--a visualization system for exploratory research and analysis. J Comput Chem 25, 1605–1612, doi:10.1002/jcc.20084 (2004).

29 Emsley, P. & Cowtan, K. Coot: model-building tools for molecular graphics. Acta Crystallogr D Biol Crystallogr 60, 2126–2132, doi:10.1107/S0907444904019158 (2004).

30 Afonine, P. V. et al. Real-space refinement in PHENIX for cryo-EM and crystallography. *Acta crystallographica. Section D*, Structural biology 74, 531–544, doi:10.1107/S2059798318006551 (2018).

31 Pettersen, E. F. et al. UCSF ChimeraX: Structure visualization for researchers, educators, and developers. Protein Sci 30, 70–82, doi:10.1002/pro.3943 (2021).

32 Jo, S., Kim, T., Iyer, V. G. & Im, W. CHARMM-GUI: a web-based graphical user interface for CHARMM. J Comput Chem 29, 1859–1865, doi:10.1002/jcc.20945 (2008).

33 Casares, D., Escriba, P. V. & Rossello, C. A. Membrane Lipid Composition: Effect on Membrane and Organelle Structure, Function and Compartmentalization and Therapeutic Avenues. Int J Mol Sci 20, doi:10.3390/ijms20092167 (2019).

34 Huang, J. et al. CHARMM36m: an improved force field for folded and intrinsically disordered proteins. Nature methods 14, 71–73, doi:10.1038/nmeth.4067 (2017).

35 Kohnke, B., Kutzner, C. & Grubmuller, H. A GPU-Accelerated Fast Multipole Method for GROMACS: Performance and Accuracy. J Chem Theory Comput 16, 6938–6949, doi:10.1021/acs.jctc.0c00744 (2020).

